# Orchid losers and winners after fire in West Australian urban bushland: A response continuum deeply integrated with other traits

**DOI:** 10.1101/2024.10.28.620558

**Authors:** Mark C. Brundrett

## Abstract

**Context:** The Southwest Australia Floristic Region (SWAFR) is a global hotspot for plant taxonomic and functional diversity with 450 highly specialised orchids facing threats from habitat loss, grazing, weeds, fire and climate change.

**Aims:** To develop fire history maps and effective monitoring tools to compare fire ecology of orchids with diverse ecological traits from an isolated urban banksia and eucalypt woodland.

**Methods:** A 72-year fire history map was intersected with >1000 orchid locations and 4.5 km of transects to measure effects on abundance and diversity. Orchid survival, size, flowering, pollination and germination were measured post-fire relative to areas unburnt for >22 years.

**Key results:** There were 58 overlapping major fires (1972-2024), averaging 8.7% of the 63-ha area annually. Correlating locations of 17 orchids with fire history revealed 5 fire-intolerant and 6 less productive species, while 6 benefited from fire-promoted flowering. Even the latter could be killed by unseasonal autumn fire. Fire decreased or increased pollination, with three orchids dependant on fire. Overall, fire impacts greatly out way benefits, as most orchids preferred long-unburnt areas, with maximum diversity and abundance three decades or more post-fire. Paradoxically, *Pyrorchis nigricans* required fire to flower, but only thrived in long-unburnt areas.

**Conclusions:** Orchids had diverse fire outcomes from catastrophic to beneficial summarised by fire response indexes (FRI) and fire age safe thresholds (FAST). This continuum was correlated with tuber depth, clonality, dispersion and lifespans of orchids, so is deeply integrated with their biology and ecology.

**Implications:** Research in an urban nature reserve provided essential tools for sustainable management of orchids relevant to rare species, such as fire history maps, FRI and FAST. Many SWAFR orchids prefer long unburnt areas, are intolerant to fire, or can be harmed by aseasonal fires. Thus, fire must be carefully managed in their habitats.

**Data Statement:** All data are provided as tables and figures or supplemental tables

## Introduction

The Southwest Floristic Region of Western Australia (SWAFR) is a global diversity hotspot with exceptional plant species richness and endemism (Myers et al. 2000, Hopper & Gioia 2004). Plant diversity in this region is linked to long-term climate stability, complex landscapes and extremely infertile soils (Lambers et al. 2010, Groom & Lamont 2015). Plants in this region have evolved exceptionally complex traits, especially for fire, pollination, and nutrition, with 90% having one or more fire survival adaptations (Brundrett 2021). This makes the region a living laboratory for studying fire trait evolution and its ecological consequences. Despite their scarcity in the arid interior, the SWAFR has an exceptional diversity of terrestrial orchids (450 taxa, https://florabase.dbca.wa.gov.au). Orchids in this region face substantial threats such as fire, weeds, grazing, excessive visitors to fragile habitats, and increasing impacts of drying and warming climate (Duncan et al. 2005, Brundrett 2014, Martín-Forés et al. 2022, Doherty et al. 2024). They also lack long-term persistent seed banks, the key to fire recovery for many SWAFR plants (70% of the flora - Brundrett 2021). However, orchid biology also provides advantages such as wind dispersed seeds and summer drought avoidance by dormancy. Biological factors that influence orchid population sustainability include very specific pollinators attracted by visual (44%) or sexual (38%) deception (Brundrett et al. 2024) and extreme nutritional dependencies on mycorrhizal fungi, since they have very few roots or lack them entirely (Ramsay et al. 1986; Brundrett 2014).

Habitats where orchids grow in south Australia are experiencing increasing fire frequency, severity, and substantial reduction in unburnt areas (Bradshaw et al. 2018, Doherty et al. 2024). They also face growing risks from drought, diseases and exotic animals (Hobbs & Atkins 1990, Groom & Lamont 2015, Wraith & Pickering 2019, Brundrett 2021, Gallagher et al. 2022). Existing information in Australia includes examples of strongly negative to positive associations with fire depending on the orchid species and fire intensity. Adams (2018) found substantially reduced orchid abundance and diversity (14 out of 33 species) after fuel reduction fires in Victoria, with only partial recovery after 10 years. Reiter and Pollard 2013 found *Caladenia disjuncta* was severely impacted by a bushfire and Jasinge et al. (2018) reported that spring or summer fires were less damaging than autumn to winter fires for a *Pterostylis* species. Duncan (2012) summarised the impacts of catastrophic 2009 fires in Victoria on orchids. These belonged to four main categories (a) fire killed species that were epiphytes or had shallow tubers, (b) fire sensitive species with substantial reductions in populations post-fire, (c) fire neutral species, (d) species where fire stimulated flowering and (e) fire dependant (pyrogenic) species that only flowered after fire. Cargill (2005) found the abundance of some orchids substantially increased with time since fire in WA forests. Australian terrestrial orchids include species that are pioneer, or secondary colonisers of fully disturbed habitats, or only occur in undisturbed habitats (Collins & Brundrett 2015). Some orchids benefit from intermediate levels of disturbance but are intolerant of substantial weed cover (Martín-Forés et al. 2022).

Fire regulates reproduction of SWAFR orchids, with 10 orchids that flower exclusively post-fire, 263 with flowering strongly promoted by fire, but 166 out of 450 taxa are considered to be fire sensitive (Brundrett 2021, Brundrett et al. 2024). Coates and Duncan (2009) reported reductions in the proportion of flowering plants of *Caladenia orientalis* with time since fire due to increased shrub competition, but rainfall was also an important determinant of flowering effort. However, Brown and York (2016) found the effect of fire on orchid pollination was less important than other factors. Fires also have impacts on specific pollinating insects and mycorrhizal fungi required by orchids. However, Marquart (2017) found limited consequences of fire on orchid pollinators in eucalypt woodlands of South Australia, while Phillips et al. (2024) found specific wasp pollinators and mycorrhizal fungi of *Caladenia tessellata* were still present after fires in southeast Australia. Overall, there was insufficient data on impacts or benefits of fire on Australia orchids relative to habitat types and species. These data are essential for substantiable management of orchid habitats

Banksia woodlands are threatened plant communities where it is especially important to monitor threats to biodiversity such as fire, weeds, isolation and climate change (Government of Western Australia 2000, Ramalho et al. 2014; Commonwealth of Australia 2016, Ritchie et al. 2021, Fowler et al. 2023). Banksia woodlands in the Swan Coastal Plain bioregion also have an exceptional diversity of indigenous terrestrial orchids, with over 150 species (Brundrett & George in prep.). This study occurred in an isolated urban remnant of banksia woodland on the Swan Coastal Plain in the Perth Metropolitan Region, which is a local hotpot of orchid diversity (Table 1, Fig. 1). The principal objectives were to investigate Indirect and direct impacts of altered fire frequency and severity on orchids in this isolated urban habitat. Specific objectives were to:

i. Produce fire history maps summarising spatial and chronological variations in fire impacts as tools for ecological research and site management.
ii. Use recently burnt and long unburnt areas to measure direct effects on orchid mortality, growth, flowering and seed production, as well as plant density effects.
iii. Measure indirect effects of time since fire on orchid distribution and productivity.
iv. Develop robust fire response and tolerance indexes and determine safe fire age thresholds for species as tools for orchid ecology and conservation.
v. Use these criteria along with inherent ecological and biological characteristics to assign orchids to fire response categories.

**Figure 1.**
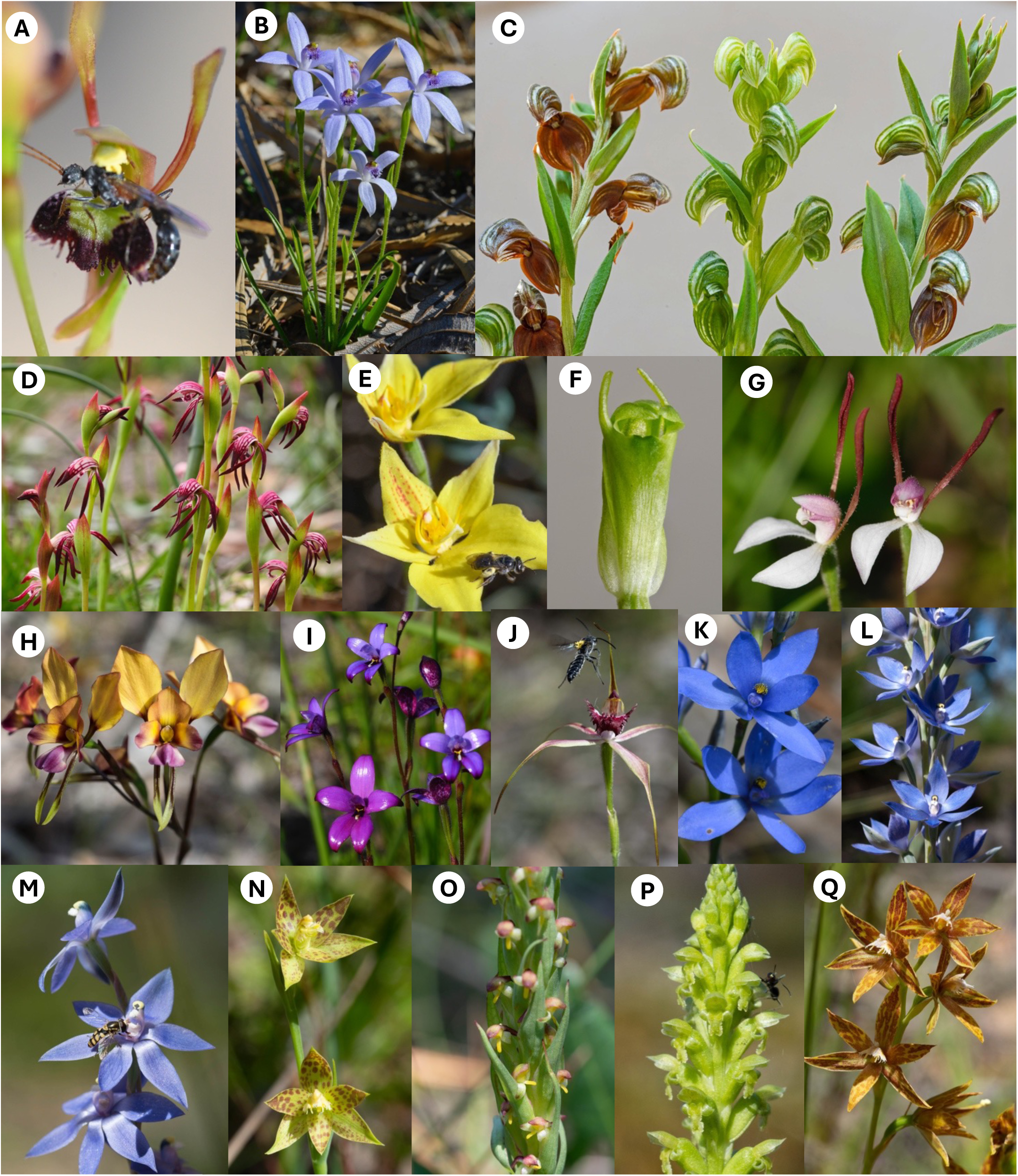
Orchid species included in study arranged by flowering sequence from autumn to early summer. **A** Leporella fimbria. **B** Pheladenia deformis. **C** Pterostylis orbiculata, **D** Pyrorchis nigricans, **E** Caladenia flava, **F** Pterostylis ectypha, **G** Leptoceras menziesii, **H** Diuris magnifica, **I** Elythranthera emarginata, **J** Caladenia arenicola, **K** Thelymitra crinita, **L** Thelymitra macrophylla, **M** Thelymitra graminea, **N** Thelymitra benthamiana, **O** Disa bracteata, **P** Microtis media, **Q** Thelymitra fuscolutea.

**Table 1.** Orchid species in this study with key biological and ecological pre-existing data.

| Orchid | Common Name | Label | Pollination syndrome | Flowering time (order) | Post-fire flowering | Flowers per spike | Capsules per flower | Organic matter in soil | Litter cover | Shade tolerance | Local abundance |
| --- | --- | --- | --- | --- | --- | --- | --- | --- | --- | --- | --- |
| *Caladenia arenicola* | Carousel spider | Ca | SD | Sep-Oct (10) | neutral | 1.4 | 0.16 | low or medium | low | low | scattered, stable |
| *Caladenia flava* | Cowslip | Cf | VD | Aug-Oct (5) | increased | 2.0 | 0.17 | high or medium | medium | medium | common most areas |
| *Disa bracteata* | South African orchid | Db | SP | Oct-Nov (160) | none | 26.3 | 0.57 | high | low | low | fairly common in some areas |
| *Diuris magnifica* | Pansy | Dm | VD | Aug-Oct (8) | neutral | 3.6 | 0.05 | various | medium | low | very common most areas |
| *Elythranthera brunonis* | Enamel | Eb | VD | Sep-Oct (9) | neutral | 1.8 | 0.15 | high or medium | medium | medium | scattered |
| *Leporella fimbria* | Hare | Lf | VD | Apr-Jun (1) | increased | 1.7 | 0.34 | low or medium | low | high | common, scattered |
| *Leptoceras menziesii* | Rabbit | Lm | SD | Aug-Sep (7) | obligate | 1.3 | 0.60 | high or medium | medium | medium | uncommon |
| *Microtis media* | Mignonette | Mm | SP | Oct-Nov (17) | none | 35.8 | 0.88 | high or medium | low | low | scattered, weedy |
| *Pheladenia deformis* | Bluebeard | Pd | VD | Jun-Aug (2) | increased | 1.0 | 0.11 | low | low | low | common, scattered |
| *Pterostylis ectypha* | Short-eared snail | Pe | SD | Aug-Sep (6) | none | 4.0 | 0.40 | high | high | high | scattered |
| *Pterostylis orbiculata* | Banded Greenhood | Po | SD | Aug-Sep (3) | neutral | 1.0 | 0.42 | high | high | high | very common |
| *Pyrorchis nigricans* | Red Beak | Pn | VD | Aug-Sep (4) | obligate | 3.5 | 0.08 | high or low | high | medium | very common |
| *Thelymitra benthamiana* | Leopard | Tb | SP | Aug-Sep (14) | no data | 13.5 | 0.03 | high or medium | medium | low | localised, increasing |
| *Thelymitra crinita* | Blue lady | Tc | VD | Aug-Sep (11) | no data | 3.9 | 0.14 | high or medium | medium | medium | localised, increasing |
| *Thelymitra fuscolutea* | Chestnut sun | Tf | VD | Aug-Sep (18) | no data | 2.4 | 0.88 | high | medium | low | localised, stable |
| *Thelymitra graminea* | Small Blue Sun | Tg | VD | Aug-Sep (13) | none | 6.7 | 0.08 | high | high | medium | localised, increasing |
| *Thelymitra macrophylla* | Blue Sun | Tm | VD | Aug-Sep (12) | rare | 6.6 | 0.06 | high | high | medium | common and increasing |
| **Main data source** |  |  | Brundrett et al. 2024 | Brundrett 2019 | Brundrett 2021 | Brundrett 2019 | Brundrett 2019 | Brundrett 2014 | Brundrett 2014 | Brundrett 2014 | Brundrett 2019 |

Pollination Syndromes: SD = sexual deception, VD = visual deception, SP = self-pollination

This research is part of a comprehensive study of all orchids present in an urban nature reserve (Brundrett 2019). Related ongoing projects are investigating variations in the reproduction and spread of orchids and the impacts of climate extremes.

## Methods

### 1. Study site and species

This study occurred in Warwick Bushland, an urban reserve of 63 hectares, located in the City of Joondalup in Perth in Western Australia (Fig. 2A). This is a Bush Forever Site protected due its size, biodiversity and vegetation condition (Government of Western Australia 2000). This is an exceptional study site due to its known fire history and large areas of very good or excellent condition vegetation. There are 215 native flora species, including 35 orchids of which 12 are locally rare in Perth (https://friendsofwarwickbushland.com accessed 18-4-2024). Vegetation is dominated by *Banksia* species with a scattered overstory of eucalypts. Important trees include *Banksia attenuata*, *B. menziesii* and *Eucalyptus marginata*, with scattered *Allocasuarina fraseriana* and *Eucalyptus gomphocephala*. It is in the Karrakatta – Central and South Vegetation Complex within the Spearwood Dune system, which is around 40,000 years old and has highly infertile pale yellow/grey sands (https://catalogue.data.wa.gov.au/dataset/perth-plant-communities-reference-sites/resource/10bedb8e-9db3-484d-9eca-2044c965cb9a).

**Figure 2.**
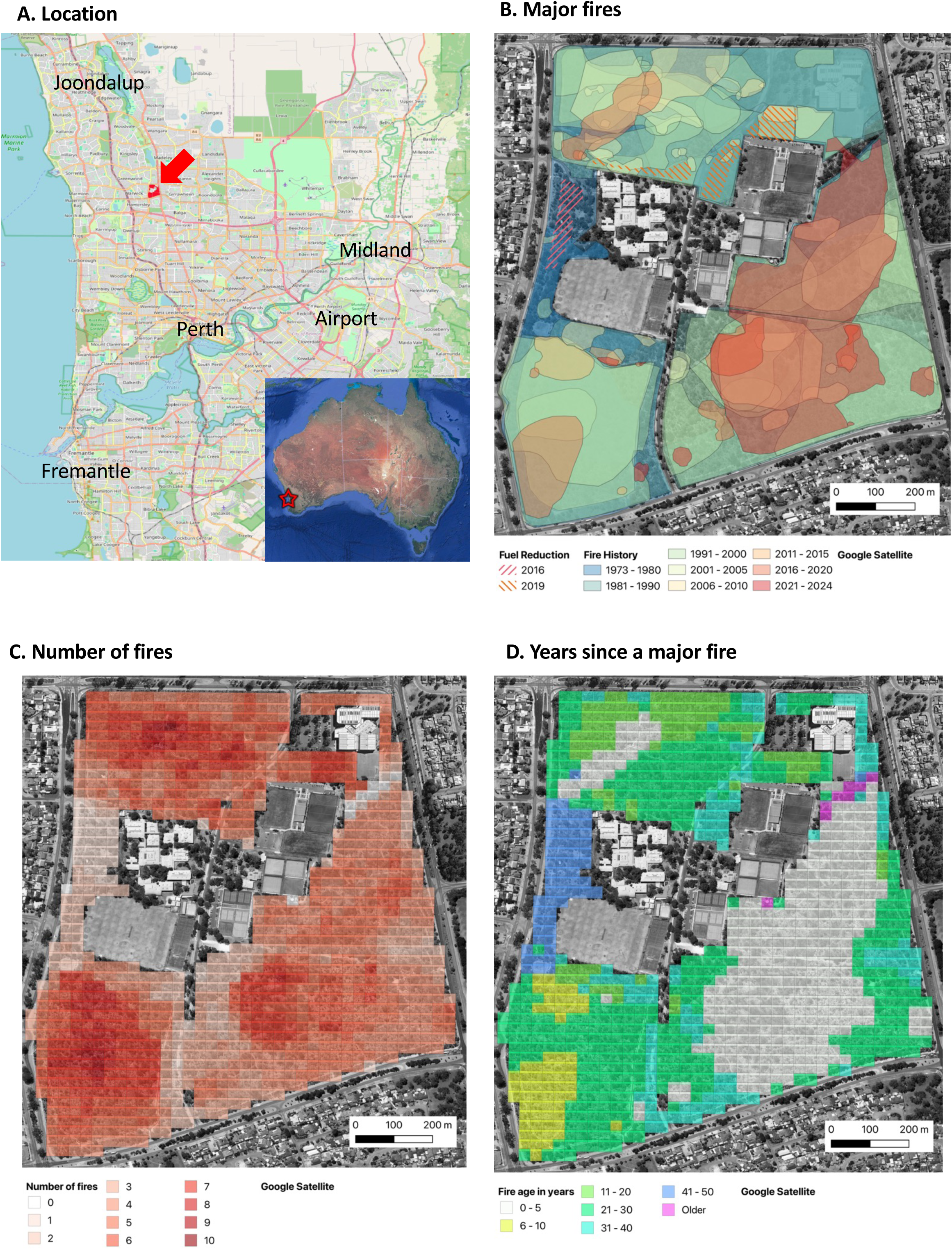
**A**. Location of Warwick Bushland. **B.** All major fires and fuel reduction burns in Warwick Bushland between 1973 and 2023. Colour shading is from the most recent in red to oldest in blue. **C**. The total number of fires in each 25 m grid square. **D**. Time since the most recent fire in each 25 m grid square. Background maps from Google.

### 2. Fire history

Data presented in Figures 2 and 3 extends a 1952-2005 fire map made by manually tracing fire scars from high resolution 300 mm^2^ contact prints of aerial photographs housed in the former WA Department of Land Administration (Clarke & Brundrett 2005). For fires post 2000, fire scars visible on digital imagery were digitised into polygons using Google Earth Pro (accessed 26-4-2024) or Nearmap imagery (https://www.nearmap.com/au/en – accessed 25-4-2024) and combined into layers using QGIS (3.34, Prizren). Only one image was available per year prior to 2002 (Google Earth Pro) or 2007 (Nearmap), when imagery become more frequent. Additional information about the dates and sizes of fires was from observations and photographs, direct mapping and maps provided by land managers (the City of Joondalup). Small fires impacting areas <10 m wide are omitted from maps but were sometimes common due to arson. Mapped major fires were primarily visible due to tree canopy loss so their polygons may be smaller than actual fire boundaries.

**Figure 3.**
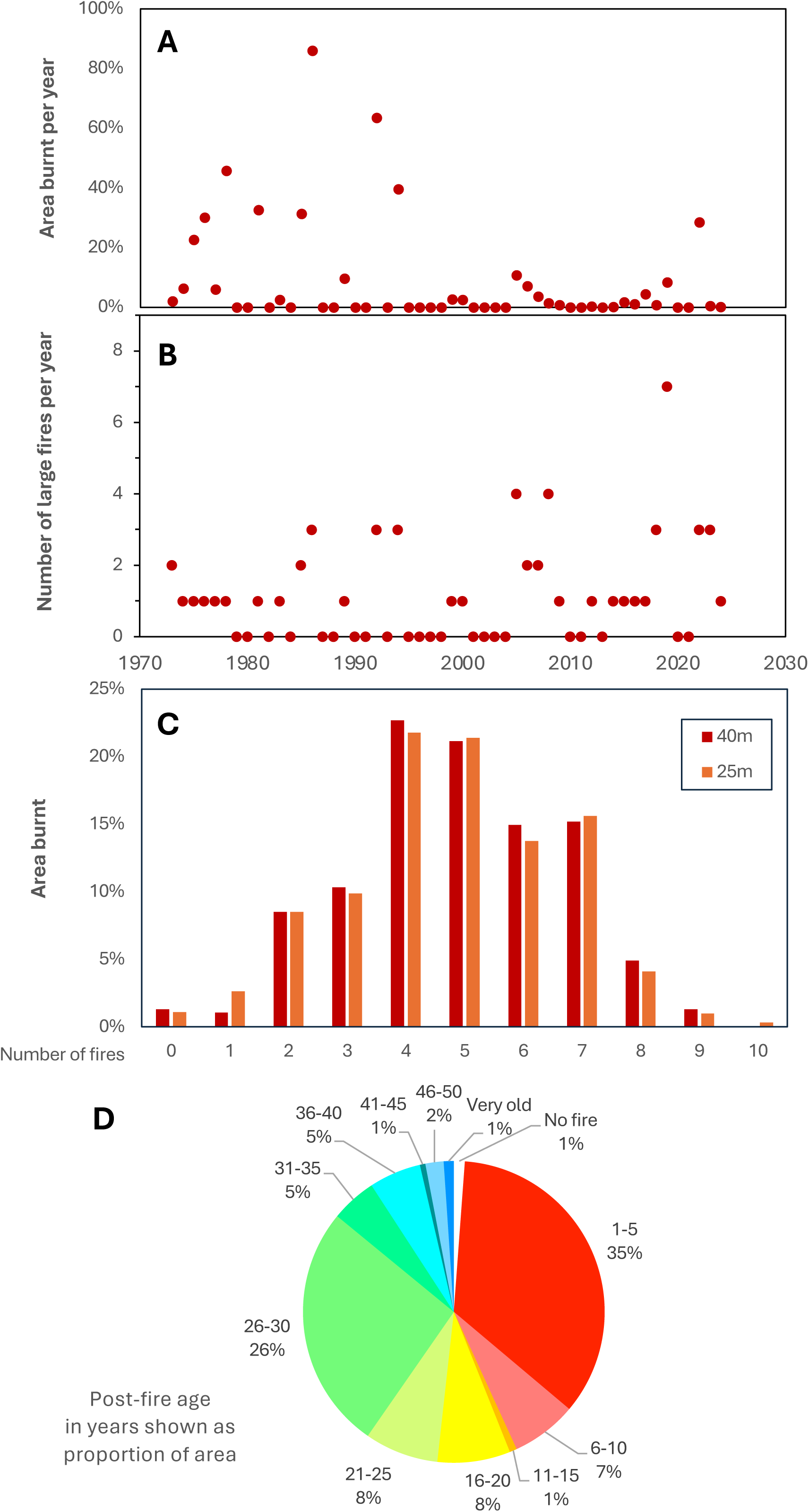
Fire history for Warwick Bushland summarising data in Figure 2. Total major fire area (**A**) or frequency (**B**) by year. Fire frequency (**C**) or age (**D**) by area.

The age of fires is defined here as the year where major fire damage was visible on imagery, so they sometimes occurred late in the previous year. These major fires usually occurred in late spring, summer, or early autumn (November - April) and had very severe impacts, resulting in complete loss of leaves and branches from understory plants and trees across most of the burn areas.

### 3. Survey methods

Comprehensive monitoring of the abundance, flowering and pollination of orchids in the study area began in 2015 and increased from 2018 onwards (Brundrett 2019). Fire studies began after a very large fire in 2022 enveloped 1/3 of the study site (18 ha). This fire impacted 200 orchid groups with 3 to 7 years of existing data, that were paired with 335 long unburnt groups for a total of ∼3000 flowering plants (Tables 2, S1). Monitoring continued for 3 years to confirm that orchids were lost to fire (some may have been grazed or dormant). Orchids missing for three continuous years are unlikely to be alive (Tremblay et al. 2009, Hutchings 2010, Brundrett 2016). Observations designated as “post-fire” occurred within a year of a fire and long unburnt areas had over 22 years since the last fire (post 1999). Several fuel-reduction burns lit in autumn or winter reduced litter and understory vegetation but had limited tree canopy impacts (Fig. 2B). These were less intense and more patchy than major fires and avoided most areas where orchids were common, so were excluded from analyses, except for survival data from a fire in June 2019 (1.3 ha).

In 2024 orchids were recorded along 30 transects (10 m wide) to increase uniformity of coverage relative to fire age (Table S2). This also ensured that spatial data included the full range of habitat conditions, from relatively weedy to almost pristine habitats. There were more transects in unburnt areas (17) than areas burnt in 2022 (13), but their total length was similar (2.3 vs. 2.4 km). Each transect was revisited three times in winter, early spring and late spring.

Orchids were counted as individual plants or recorded within groups - defined as aggregations of plants separated from others of the same species by several times their group diameter. Groups of some orchids were of clonal origin, but seeding orchids were also often aggregated. Most groups were >10 m wide, except for *Thelymitra macrophylla*, *P. nigricans* and *Leporella fimbria* which had colonies up to 30 m across. For consistent monitoring of the same orchid plants, locations were identified using georeferenced photos showing surrounding features and markers. These were recorded with a mobile phone using the SW Maps Android phone app (3-10 m accuracy). Orchids were visited multiple times per year to count flowering plants, non-flowering plants, seedlings, flowers per spike and capsules (all had a single spike per flowering plant). Fire impacts on plant height were also measured in 2022. Baseline data was provided by 6 to 11 years of monitoring the same orchid groups at Warwick Bushland (Brundrett 2019) totalling over 30,000 observations of plants (20,000 spikes, 140,000 flowers and 40,000 capsules - Table S1).

### 4. Fire response and trait data

Short-term fire impacts were measured using data from the same orchid groups for 3 or more years before and after a fire (Table 2). These observations primarily concern a large fire in 2022 and two smaller fires in 2023 and 2024 and included representatives of common orchids, and all individuals found for less common species (Tables S1, S3). Fire responses or traits measured include seed production, seedlings, group sizes, clonality, aggregation and dispersion, as well as tuber size, depth and abundance, with details of methods provided in Table 4. Seed production was the proportion of capsules per flower that reach full maturity, which may be less than pollination rates due to seed abortion and grazing (three species self-pollinate). *Ex situ* seed baiting (Brundrett et al. 2003) confirmed a high degree of seed viability for all species (MCB unpublished data). Plants were designated as clonal if they produced more than one tuber per plant per year. Their daughter plants are unconnected and may include different genets, so were counted as individuals. Separate indexes were calculated for clonality, aggregation and dispersion, recolonisation potential and lifespan from demographic data (Tables S1-S4), as explained in Table 4.

**Table 2.**
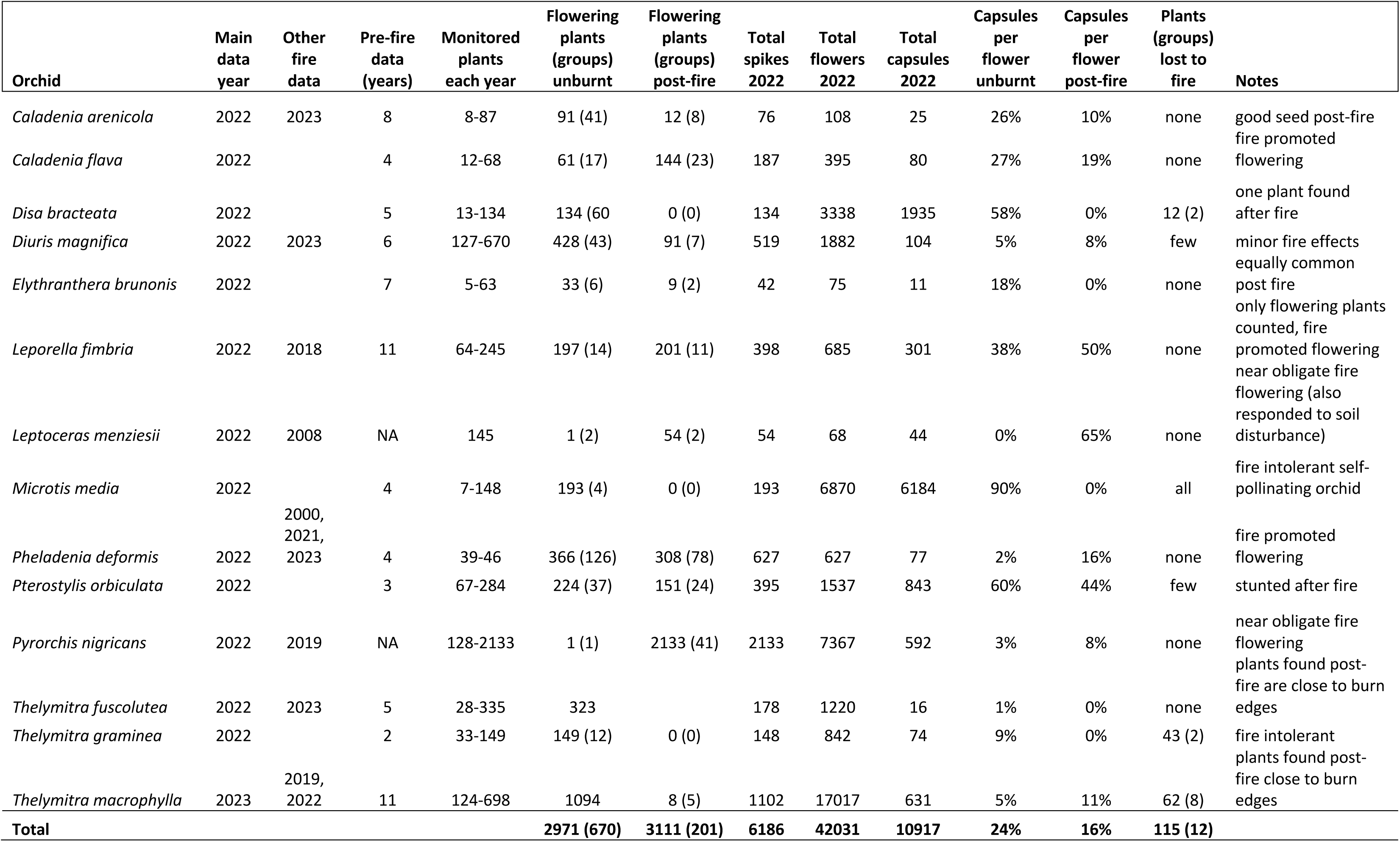
Direct effects of fire on flowering, seed set and survival of 17 orchid species.

**Table 3.**
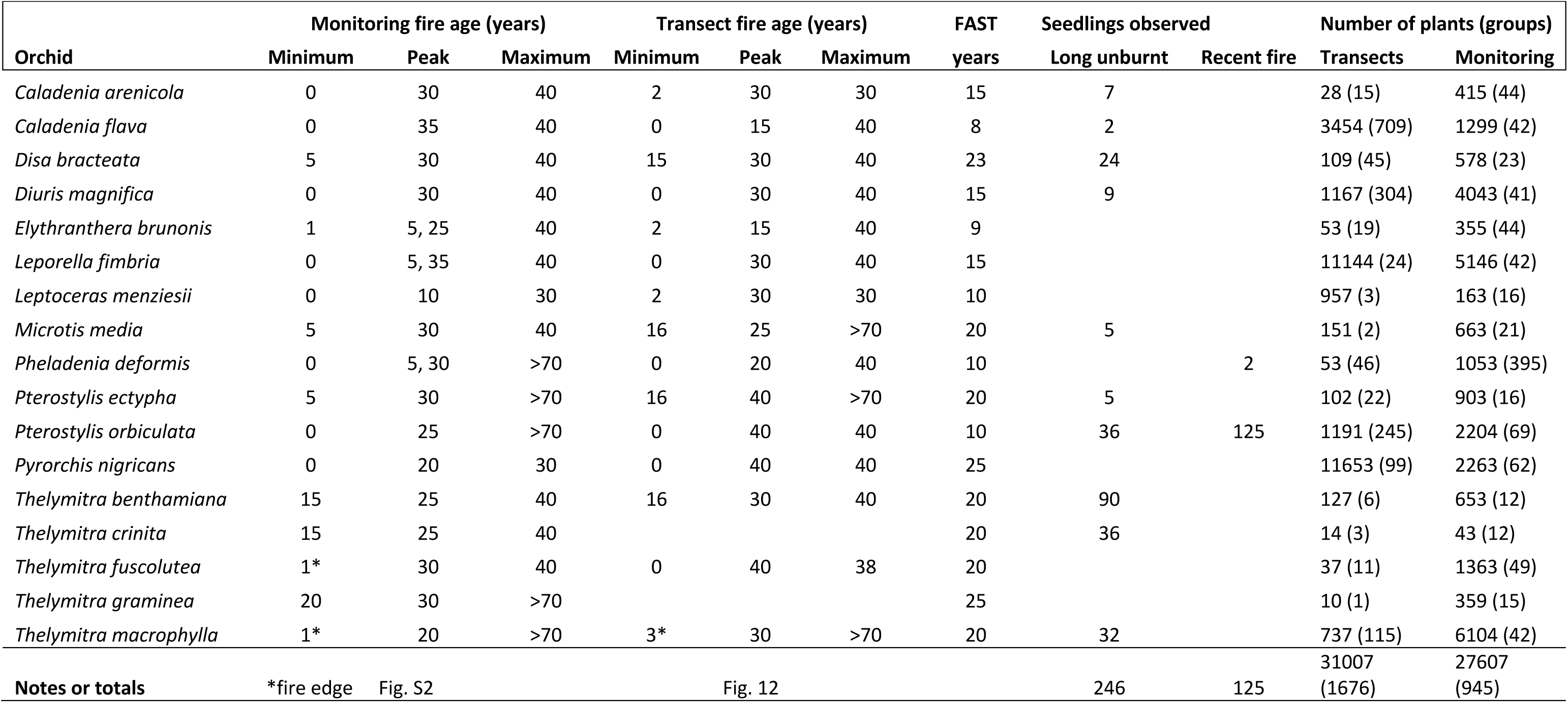
Indirect effects of fire on abundance of 17 orchid species with numbers of plants and groups counted (FAST is Fire Age Safe Threshold).

**Table 4.** List of traits linked to fire responses and indices used in Table 5 with explanations and data sources.

|  | Trait or response category | Data sources and methods | Data type | Data sources |
| --- | --- | --- | --- | --- |
| **A** | **Fire responses** |  |  |  |
| 1 | Post-fire mortality | Counts of plants and groups absent for three years after fire | C, N | Figs. 5C, 6 |
| 2 | Abundance relative to time since fire | Group sizes and locations joined with map of all fires | N | Figs. 11-13 |
| 3 | Abundance after recent fires | Group sizes joined with map of all fires or recent fires (post 1999) | C, N | Figs. S3, S4 |
| 4 | Susceptibility to unseasonal fires | Counts of orchids lost to fire in late autumn (when shoots are emerging) | C, N | Table 2, Fig. 4D, E |
| 5 | Post-fire flowering promotion or reduction | Ration of post-fire relative to long unburnt areas on the same year | C, N | Fig. 7B, C, Fig. 11 |
| 6 | Plant size after fire vs long unburnt areas | as above | C, N | Fig. 7A |
| 7 | Pollination and seed production fire response | as above | C, N | Table 1 |
| 8 | Fire intolerance | Consistent absence from recently burnt areas | C | Table 3 |
| 9 | Fire response summary | Sum of all fire traits over 6 intervals from 18 to -18, 0 = neutral | C | Table 5A |
| 10 | Fire age safe threshold (FAST) | Median of minimum and peak habitat age post-fire from abundance in monitored areas and transects | N | Table 3, Figs. 12, S2 |
| **B** | **Reproduction and ecology** |  | N |  |
| 11 | Average seed production per spike | as above by per spike (flowering plant) | N | Figs. 8, 9 |
| 12 | Availability of pollinators | Relationship between plant density and pollination rate from graphs | C | Fig. 9 |
| 13 | Seed production capacity | Long-term averages measured in unburnt areas (excludes fire-flowering species) | N | Table 1, Fig. 8 |
| 14 | Seedling recruitment | Numbers of seedlings or small young plants observed and areas where found | N | Table 3, Fig. 4 |
| 15 | Clonality | Relative abundance of in clonal groups relative to individual plants | N | Table S4 |
| 16 | Aggregation | Ratio of plants in groups to scattered individuals | N | Table S4 |
| 17 | Colonisation | Groups and seedlings found in new locations relative to totals | N | Table S4 |
| 18 | Dispersion | Ratio of numbers of orchids found by uniform (transect) sampling relative to targeted sampling to maximise diversity sampled | N | Table S4 |
| 19 | Recolonisation capacity after severe disturbance | Data from a time-course study in a bauxite mine from bare soil | C | Collins & Brundrett 1995 |
| 20 | Typical plant lifespan | Proportion of groups or plants lost during long-term monitoring of unburnt areas | N | Preliminary data |
| 21 | Tuber size and number per plant | 10 or more plants measured non-destructively | N | Table S3 |
| 22 | Tuber crown bud depth in soil | As above | N | Table S3 |

### 5. Data analysis

Impacts of fire on pollination density dependence were determined by graphing pollination outcomes against the total number of flowers in groups (Brundrett 2019). Orchid productivity was compared to local fire history using the “Join attributes by location” data analysis tool in QGIS to intersect orchid group locations or transects with fire history polygons. This produced new point layers for all fires at each orchid location as a data matrix. The age of areas without any fires was set the first year where fire data was available (1952) but is likely would be much older. To summarise fire history for the entire site area, the same QGIS tool was used to intersect a grid with fire history polygons (Fig. 2CD). Results for 25 m and 40 m grids were similar (Fig. 3C), so only the 25 m grid (1024 points) was used. Orchid locations and grid points located near fire polygon borders were manually corrected where required.

Measured fire responses were converted into categorical response indexes, extending from catastrophic impacts to major benefits converted to a unified numeric scale (-3, -2, -1, 0 1, 2, 3) as explained in Table 5A. These nine indexes were then combined to produce an overall Fire Response Index (FRI, Table 5A). Separate indices for important ecological or biological traits (Tables S1-4) were also calculated as explained in Table 4 and standardised by dividing by the largest value (x10) as a whole number (0-10).

**Table 5A.**
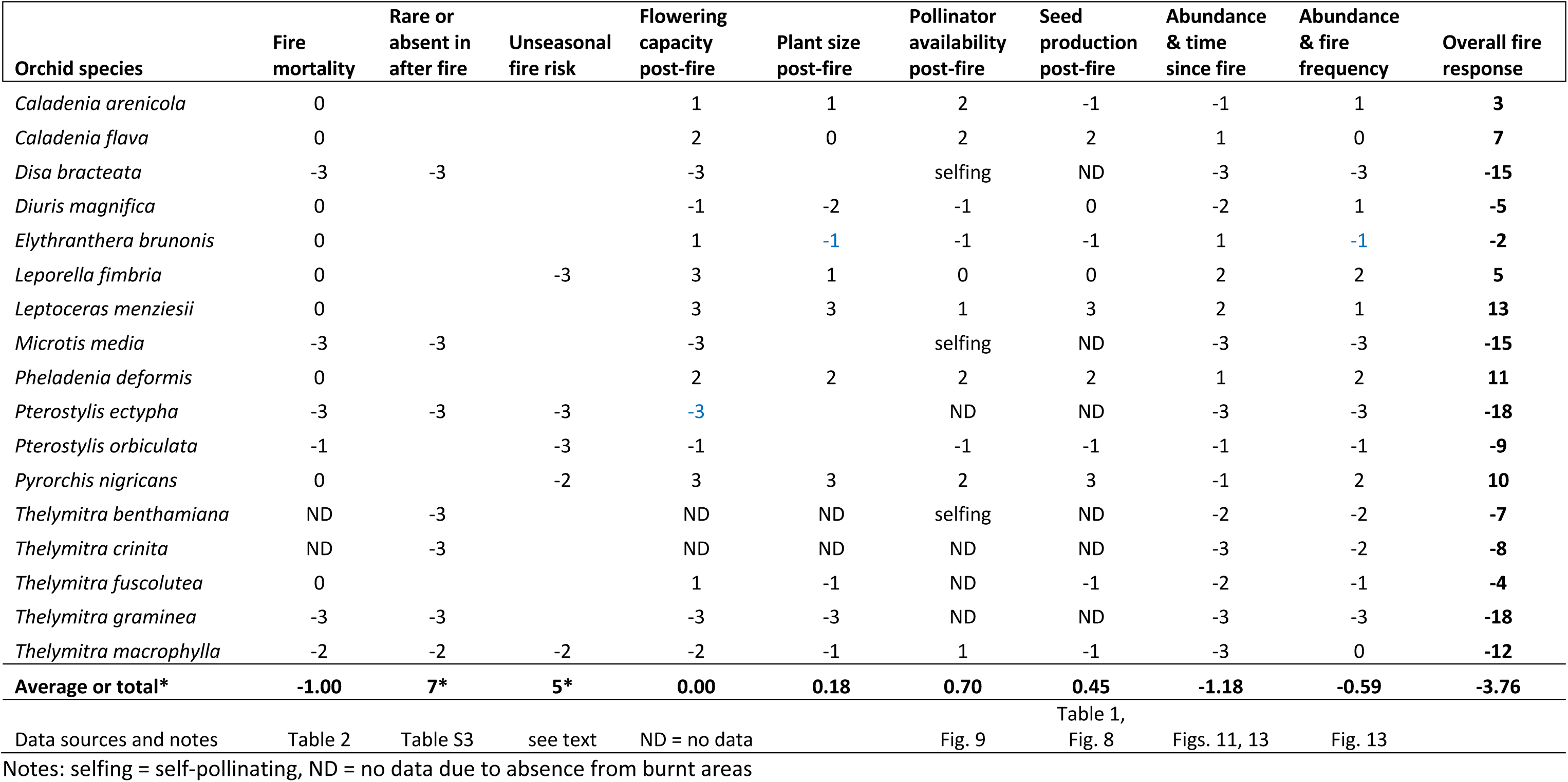
Orchid fire responses on a unified categorical scale: rare or absent (-3), greatly decreased / major impacts (-2), decreased / minor impacts (-1), neutral / negligible impacts (0), increased / minor benefits (1), greatly increased / major benefits (2), fire opportunist (3). Fire responses are based on data in Tables 1-3, as explained in Table 4.

These indices represent the capacity of orchids to spread locally or distally by clonal or seed propagation, aggregation of plants, orchid abundance trends and estimated lifespans, as well as tuber depth and size (Table S5). Typical population increases or decreases for orchid groups were determined using 3-5 years of data and divided by the average size of groups to estimate turnover rates (Table S4). Fire response data from this study was also compared with existing ecological information for each orchid (Table 1). Fire tolerances for orchid species were summarised using safe fire thresholds (FAST) and optimum fire ages, as explained in Table 4. Correlations between fire responses and traits were calculated and P values shown on graphs. Statistical comparisons of orchids between recently burnt and long unburnt areas were provided by 95% confidence intervals.

## Results and Discussion

### 1. Fire history

Major fires are mapped in Figure 2 and summarised in Figure 3. Typically, fires were of high or very high intensity (Fig. 4A, B) in summer or autumn and their sizes ranged from <100 m^2^ to most of the 63-ha total area (Fig. 3A). The entire area has been burnt multiple times since 1972, excluding a narrow zone in the northeast (Fig. 2B). Fires were generally smaller after 1995, with only the 2022 fire exceeding 10 ha, while many earlier fires consumed entire compartments bounded by tracks (Fig. 3A). This likely resulted from increased accessibility to fire vehicles (new tracks and gates), but local firefighting resources and protocols also changed (e.g. use of water bombing aircraft - Fig. 4B). All fires were caused by arson, except two managed fuel reduction fires (April 2016 - 0.7 ha and June 2019 - 1.7 ha). The latter limited tree canopy damage but had severe understory scorching (Fig. 4C). Fire polygons may include less impacted areas, especially near tracks and edges.

**Figure 4.**
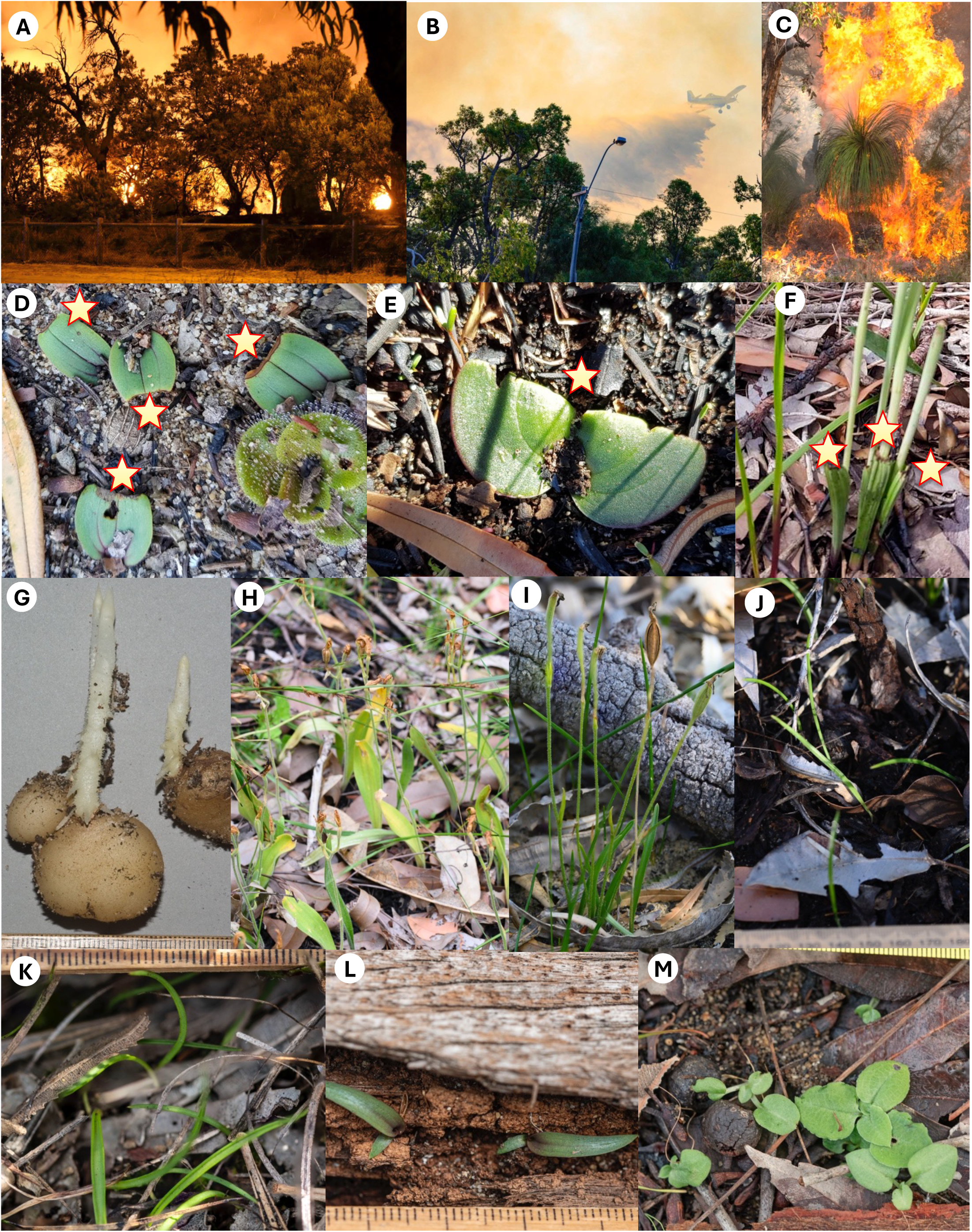
An intense summer arson fire in 2022 (**A**) with water bomber (**B**). Winter fuel reduction burn (**C**). Leaf damage (stars) to orchids by unseasonal fire in late autumn for *Leporella fimbria* (**D**) and *Pyrorchis nigricans* (**E**). **F.** Scorching of *Thelymitra macrophylla* by the fuel reduction burn (stars). **G**. *Pterostylis orbiculata* tubers sprout several months before emerging. **H**. *Caladenia flava* seed capsules after fire. **I**. *Pheladenia deformis* seed formed after fire. Seedlings of *Diuris magnifica* (**J**), *Thelymitra benthamiana* (**K**) and *Disa bracteata* on wood in a long unburnt area (**L**). **M**. *Pterostylis orbiculata* seedings three years post-fire.

On average, 1.1 severe fires occurred annually after 1972, and 5.6 ha was burnt each year (Fig. 3A). In total, 58 major fires covered a cumulative area of 289 ha (4.6 x the reserve size). There was a major fire on 30 out of the 52 years (often more than one) and a maximum of seven per year (Fig. 3B). Images were available from 1952 onwards, but fires only commenced after surrounding areas were developed for housing in the 1970s. Overall, the size of fires has decreased over time, but their frequency has increased (Fig. 3A, B). Due to the complexity of fire patterns, a 25 m grid was used to summarise average fire frequency and time since fire on maps (Figs. 2B, C, 3C, D). Fire frequency had a normal distribution with an average of 5 fires per location and a maximum of 10 (Fig. 3C). The largest areas were 1-5 years (35%) or 26-30 years post-fire (26%). This bimodal fire age distribution provided similar areas of recently burnt or long unburnt habitats for comparison (Fig. 3D).

### 2. Direct impacts on orchid survival

Figure 5 shows the distributions of orchids in Warwick Bushland relative to recent fire history and indicates where some were lost, primarily in 2022 (Fig. 5C). These loses are significant because most orchid species in the study area are stable or expanding in numbers (Brundrett 2019, Table S4). *Thelymitra macrophylla* (blue sun orchid) has rapidly increased in abundance and distribution in the study area since 42 plants in three groups were discovered in 2009 (Brundrett 2019). Six *T. macrophylla* groups were lost to fire in 2022 and still missing after 3 years (Fig. 5C). One *T. macrophylla* group with 44 flowering plants pre-fire, was much smaller after a fire in 2019 and completely gone after a second severe fire in the same area in 2022 (Fig. 6). Three groups survived a smaller fire in 2023 (Fig. 5C), but two of these were absent in 2024, when an additional surviving plant emerged. Only 11% of of *T. macrophylla* plants survived fire and these were located near fire boundaries where intensity was lower. None of the 31 monitored groups in unburnt areas were lost, some expanded substantially (up to 436 plants), and new groups were found each year (Fig. 6, Table S1). Only 10 *T. macrophylla* plants were found on transects in burnt areas, while 728 were found in long unburnt areas (Fig. 5D). The smaller blue sun orchid (*T. graminea*) also did not survive fire (43 plants in 2 areas lost). These orchids are vulnerable to autumn fires, as explained above.

**Figure 5.**
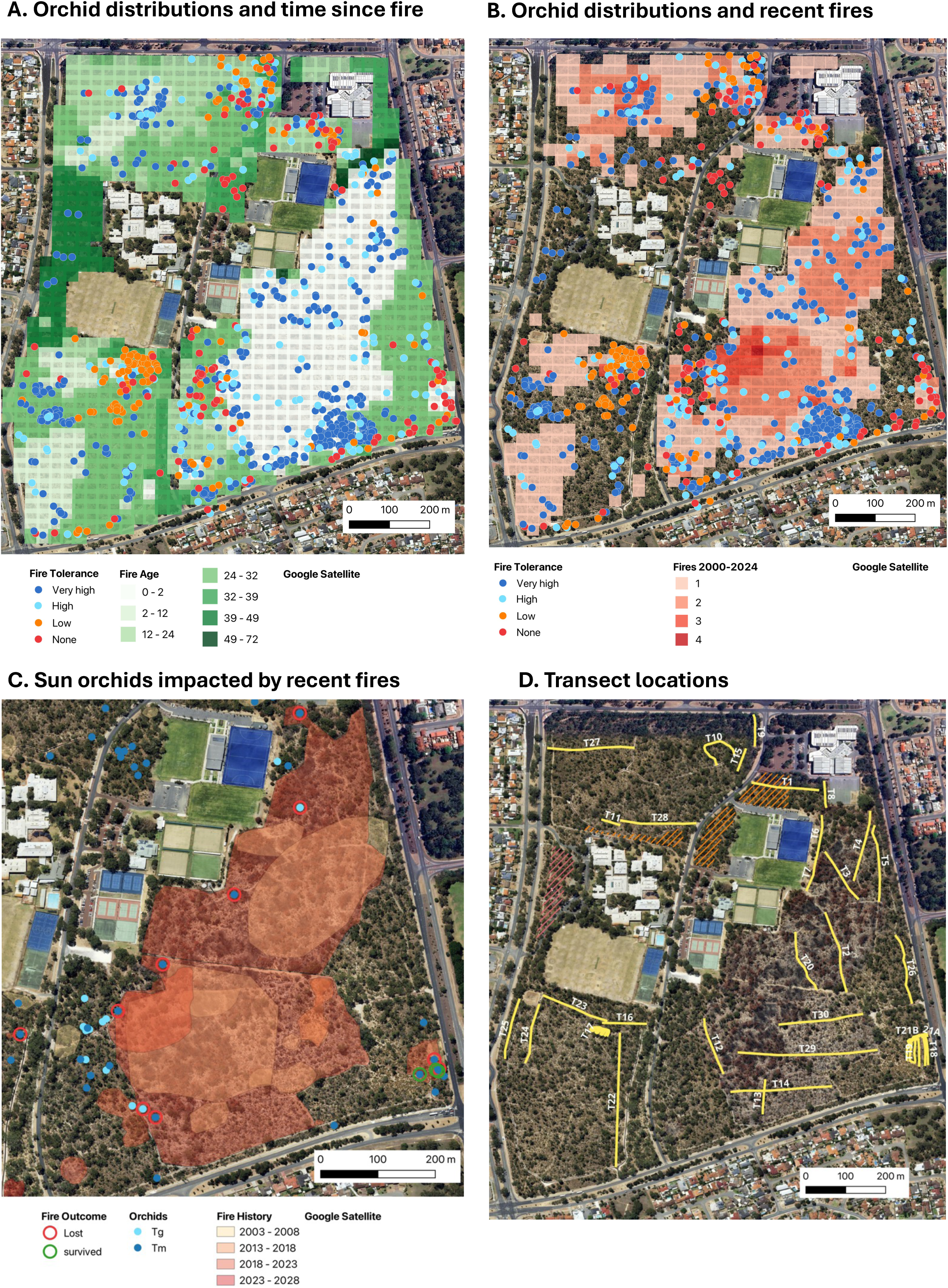
The distribution and survival of ∼1000 monitored groups of 17 orchids relative to (**A**) fire age and (**B**) recent fire frequency (2000 -2023). **C.** The distribution of sun orchids (*Thelymitra* spp.) relative to recent fires. Locations where sun orchids were killed or survived fires in 2022 or 2023 are indicated by circles (Tm = *Thelymitra macrophylla*, Tg = *T. graminea*). **D**. Transects used to measure orchid abundance and diversity relative to fire history.

**Figure 6.**
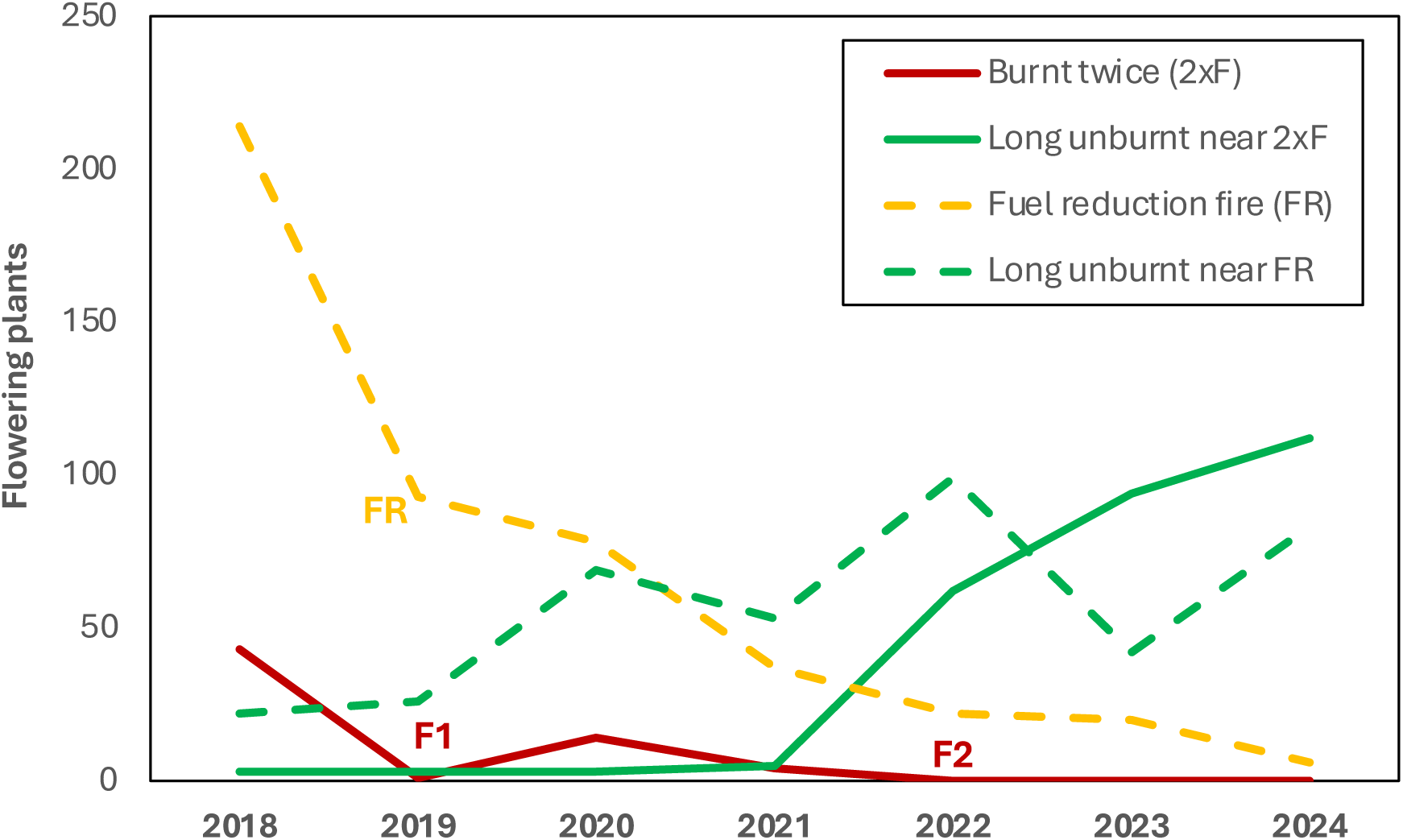
Changes over time in abundance of flowering *Thelymitra macrophylla* groups showing impacts of multiple very hot summer fires (F1, F2) and a cooler winter fuel reduction burn (FR) relative to unburnt groups of similar age nearby.

A group of over 350 *T. macrophylla* plants was impacted by a fuel reduction burn in June 2019 (Fig. 6). This burn was managed to limit damage to tree canopies, but ground level intensity was very high due to the highly flammable leaves of *Xanthorrhoea priesii* (Fig. 4C). This winter fire resulted in substantial damage to growing *T. macrophylla* leaves, but spikes continued to develop (Fig. 4F). However, impacts to this group are ongoing after five years with further declines in numbers each year, while adjacent unburnt groups expanded (Fig. 6).

Some orchids were consistently absent from recently burnt areas but common elsewhere. These include *Disa bracteata* (South African orchid) and *Microtis media* (mignonette orchid) which prefer relatively open areas near tracks and reserve boundaries (Fig. 1 O,P)*. Disa bracteata* is an especially good indicator of recent fires, with some groups bisected by fire boundaries. This orchid was also lost to fire at another site 35 km away (Brundrett & Longman 2016). These disturbance tolerant orchids have relatively shallow tubers, are short-lived and reproduce by seed (Table 5B). *Pterostylis ectypha* (a snail orchid) is also fire intolerant (Fig. 1F). Ramsay et al. (1986) found orchids in this groups had very shallow tubers, located just under the litter layer. Most fire intolerant orchids produce abundant seed (*D. bracteata* and *M. media* self-pollinate) so are expected to recolonise burnt areas as conditions become suitable.

**Table 5B.**
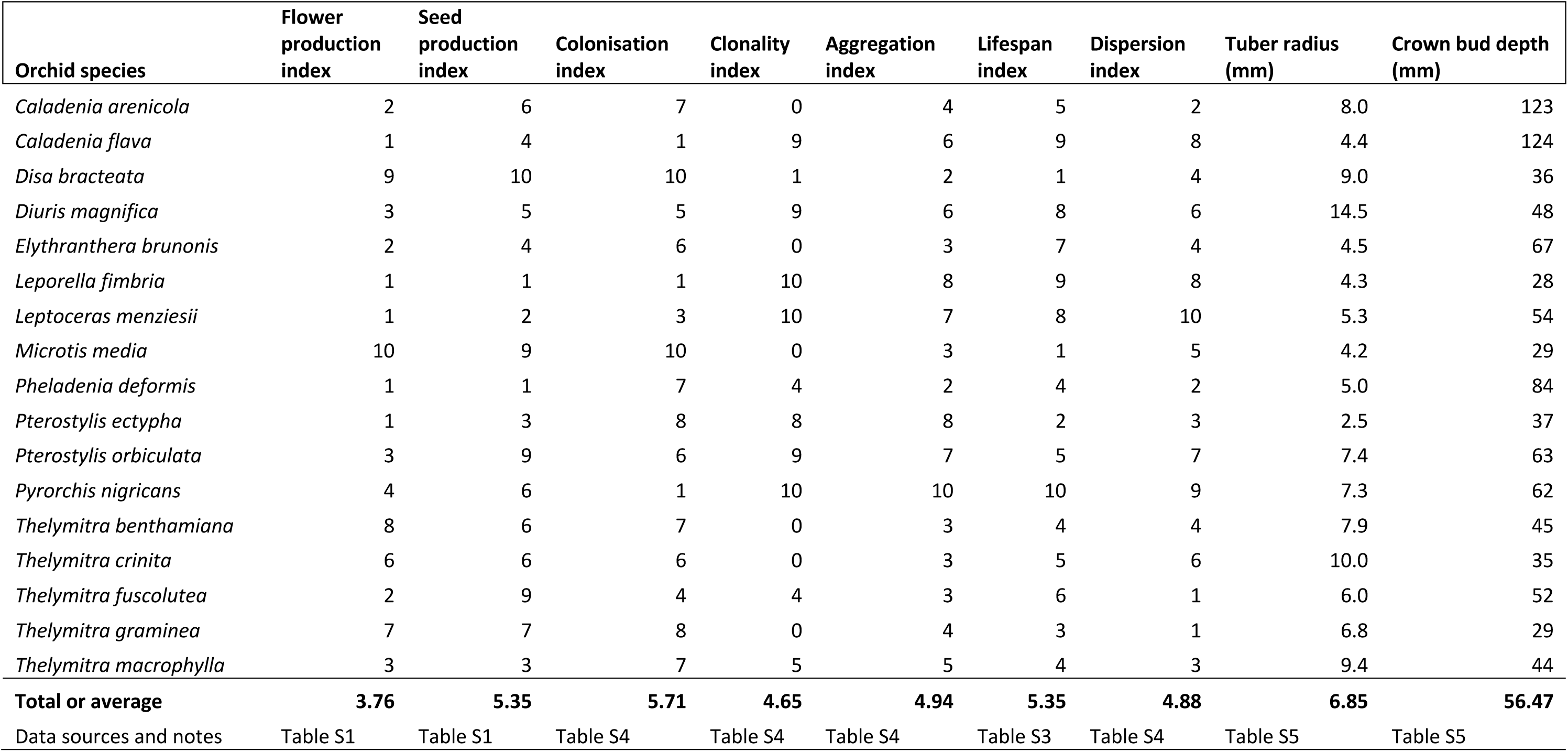
Orchid trait indexes (B) converted to a common scale (0-10) based on Tables S1-S5 as explained in Table 4.

*Leporella fimbria* (hare orchid) and *Pyrorchis nigricans* (red beaks) are normally highly resilient to fire, but an unseasonal hot fire in late autumn killed many plants in a group (May 2018). These orchids form shoots just below the soil surface in autumn and do not initiate new tubers until winter (Fig. 4G). Plants with partly scorched leaves (Fig. 4D, E) were found on the margin of the group, which had not recovered substantially after seven years. *Pterostylis orbiculata* (banded greenhood) was less productive in burnt areas but still common (see below). This orchid shoots in autumn (Fig. 4G) so is vulnerable to unseasonal fires, as is the case for *T. macrophylla* (Table 4).

### 3. Direct effects on orchid reproduction

Orchid flower production and pollination post-fire was compared to long unburnt areas on the same year (2022 unless stated otherwise). *Leporella fimbria* occurs across southern Australia and has small green and brown flowers (Fig. 2a) that attract flying male bull ants (*Myrmecia* spp.) using a pheromone (see https://www.youtube.com/watch?v=D1FTrJaEFhE&t=89s). Most plants (89%) of this long-lived, highly clonal species remain sterile, and its flowering can be severely impacted by autumn drought (Brundrett 2019). Flowering and seed set were moderately higher (11 %) in burnt areas, especially in large patches (Figs. 7, 8), but seasonal variations in rainfall were more important. This orchid has a strong density effect on pollination which was reduced after fire (Fig. 9A). Overall, increased seed production post-fire results from greater flower production but still shows that the pollinator (*Myrmecia* sp.) was present in severely burnt areas. Grazing resulted in loss of spikes each year (long-term average 6.2%) but was lower in burnt areas (4% vs. 9% of spikes).

**Figure 7.**
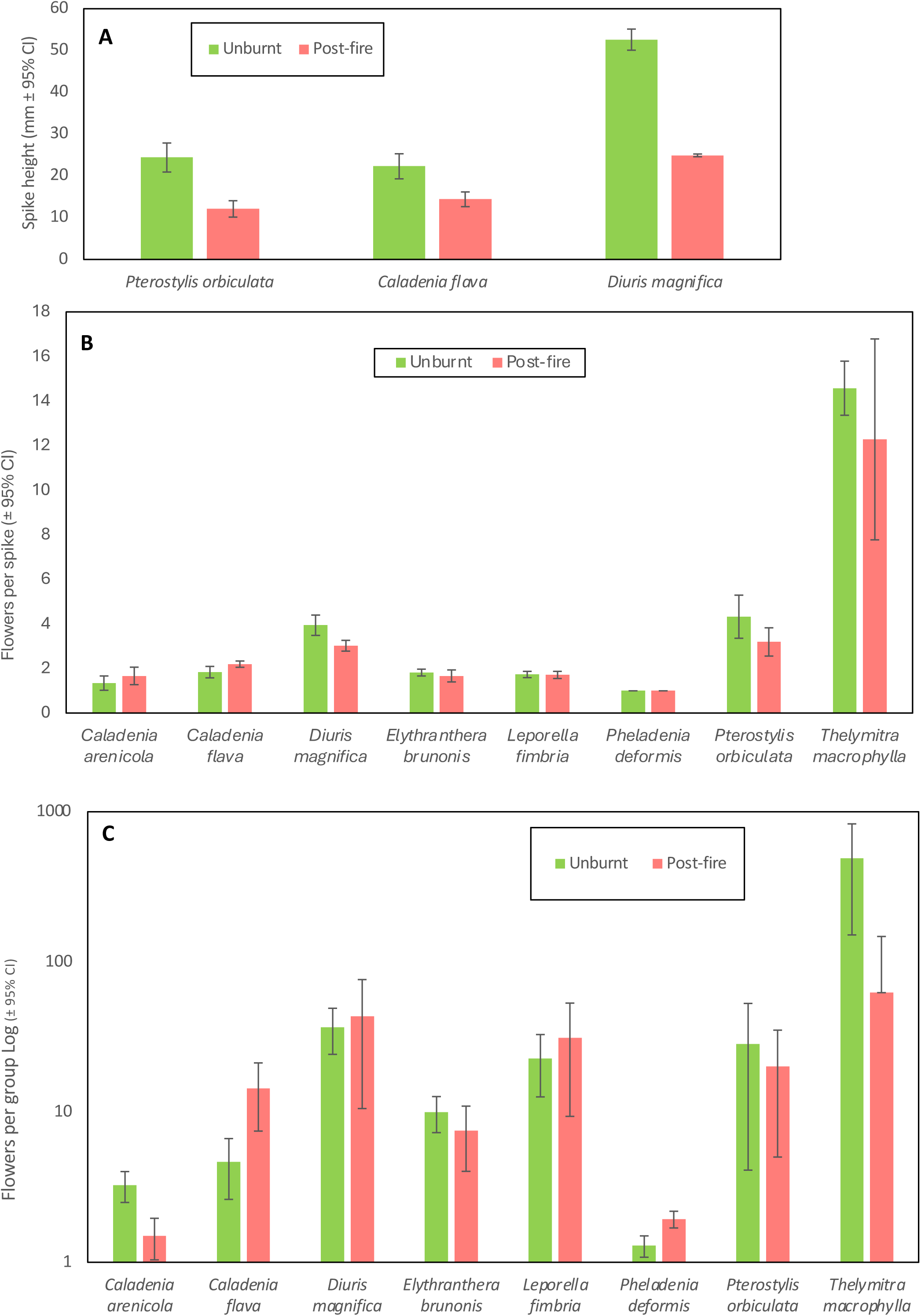
Fire effects on plant growth (**A**) and flower production per plant (**B**) or group (**C**) for common orchid species comparing long unburnt and post-fire areas.

*Pheladenia deformis* (blue beard orchid) also occurs in eastern Australia and is a close relative of *Caladenia* and *Cyanicula* (Fig. 1B). It is a relatively solitary winter-flowering orchid that can form clumps, and often occurs within 5-20 m of others due to seed dispersal or habitat preferences. Many new plants were discovered after the large 2022 fire with another geophyte not previously recorded in banksia woodland (*Wurmbea dioica*). *Pheladenia deformis* leaves were not substantially larger post-fire, but there were 1.6x more flowers per area (Fig. 7C). Pollination was very substantially increased (19x) after fire (Table 2, Figs. 4I, 8), with a moderate density effect (Fig. 9B). Outcomes differed each year and after each fire, probably due to climactic factors (Fig. 10A). Pollination was very rare in unburnt areas except in 2021. The 2022 fire produced 54% of all capsules formed over 3 years, and a very small fire produced 1/2 the 2023 capsules (Fig. 10A). This orchid is very small, so easily overlooked when not flowering. Most plants that flowered in 2022 were rediscovered as leaves only in 2023, by carefully checking the same locations. This orchid gradually recovered flowering capacity after fire over a decade (Fig. 10B). Sterile or dormant plants were also more common the second year after (Fig. 10C). Reproductive costs can reduce the likelihood of subsequent flowering in other orchids (e.g. Shefferson et al. 2020).

**Figure 8.**
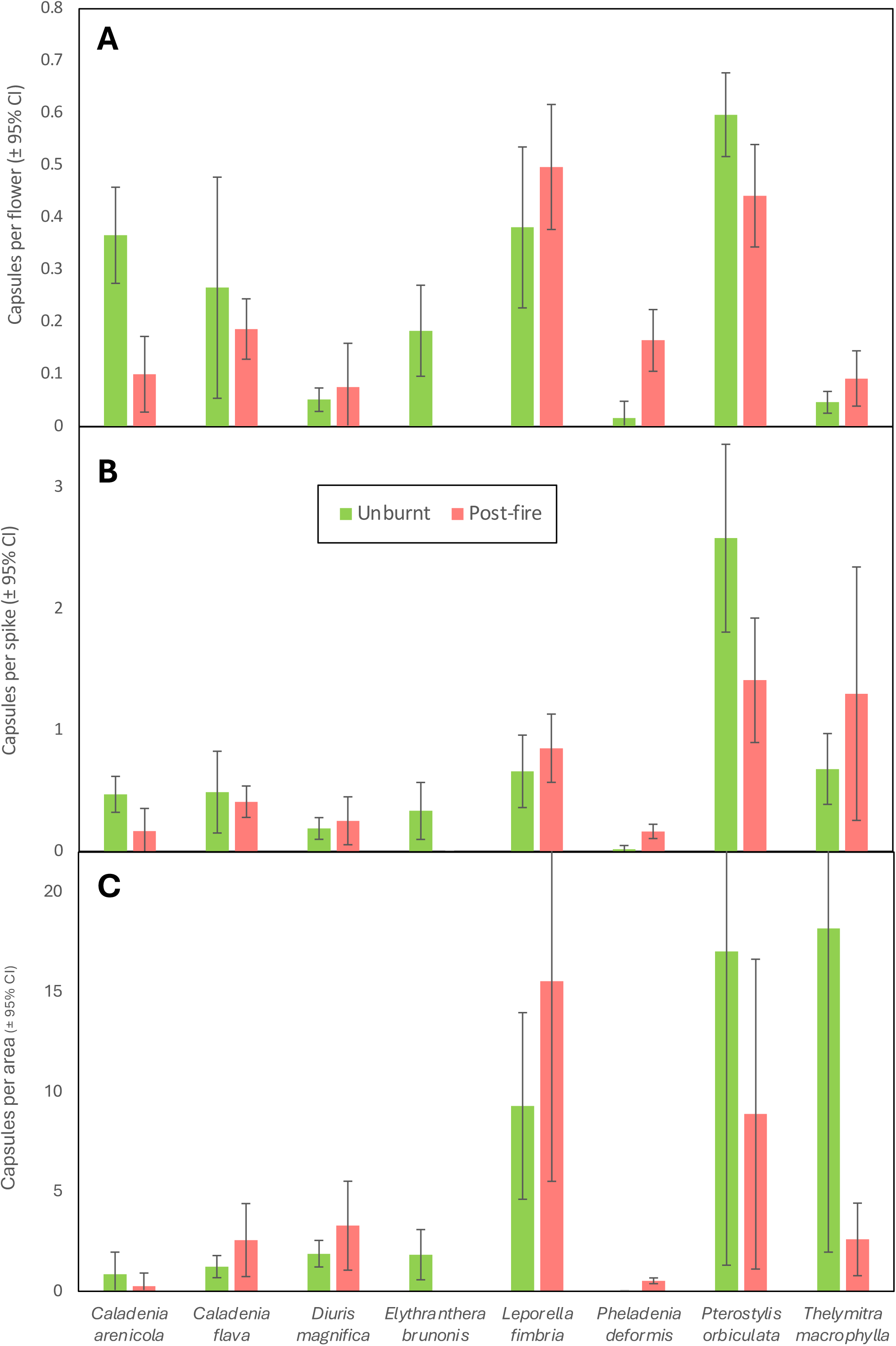
Effects of fire on overall pollination rates of eight common orchid species (see labels) summarised per flower (**A**), plant (**B**), or group (**C**).

**Figure 9.**
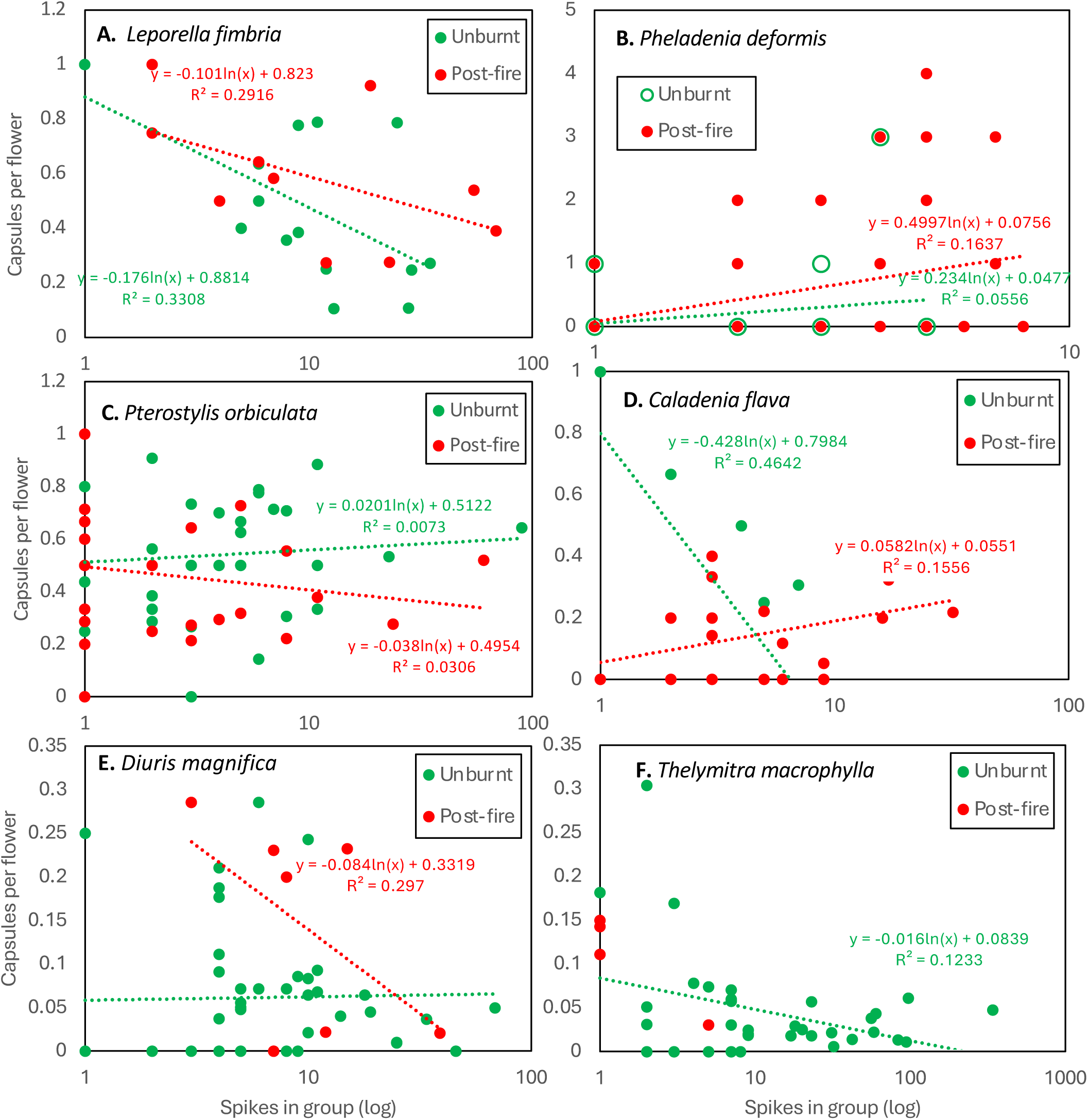
**A-F.** Effects of plant density and fire on pollination of six orchid species (see labels) in long unburnt or post-fire areas.

**Figure 10.**
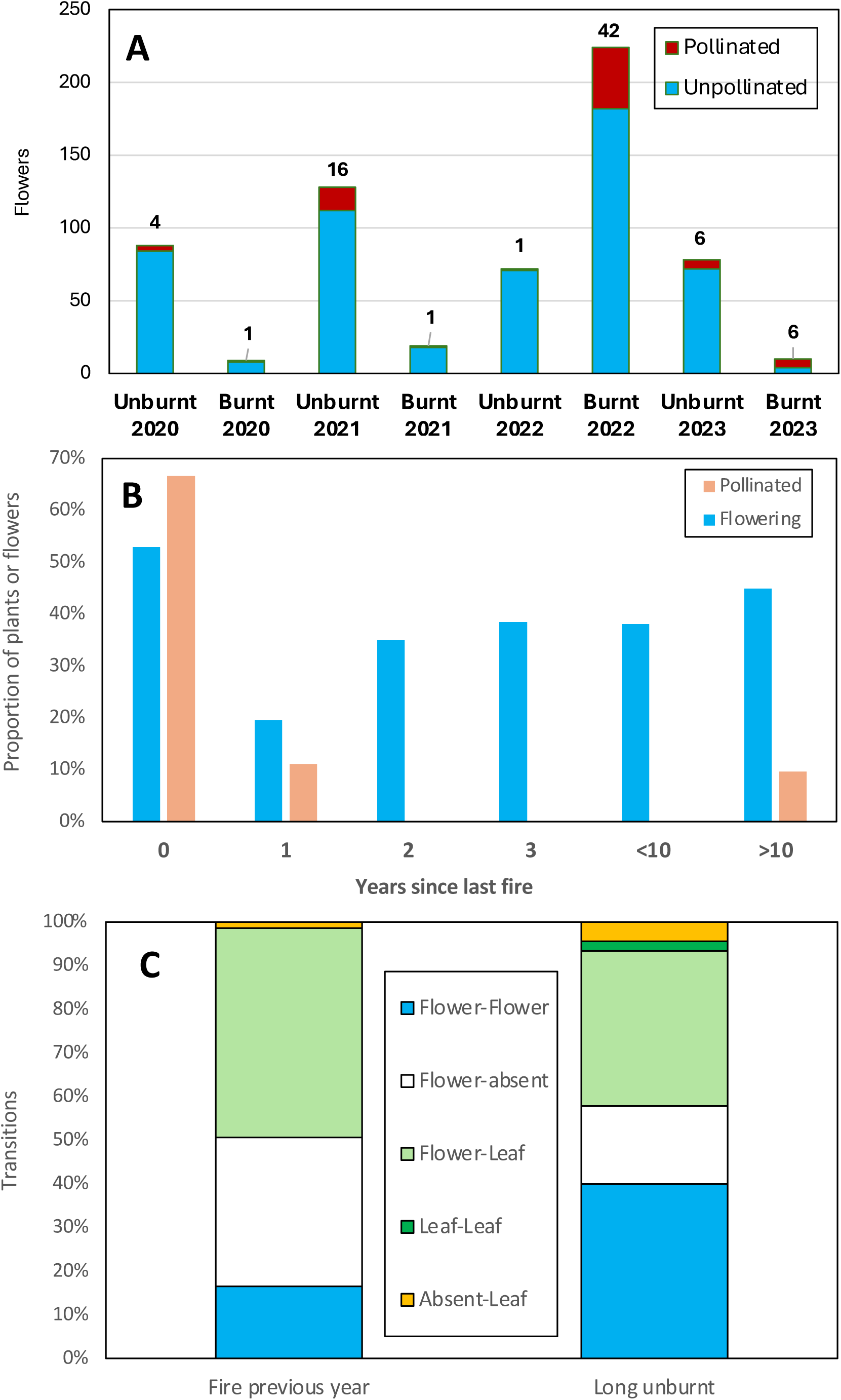
*Pheladenia deformis* pollination ecology. **A**. Pollination of burnt or unburnt plants. **B**. Changes to flowering capacity with time post fire. **C.** Transitions between vegetative, flowering or missing plants.

*Pterostylis orbiculata* (Fig. 1C) belongs to a complex of similar species that flower in winter (previously known as *Pterostylis sanguinea*). This orchid was substantially smaller after fire with fewer flowers and seeds (Figs. 7, 8). There also was a strong plant density effect on pollination suggesting there were fewer fungus gnat pollinators (Fig. 9C, https://www.youtube.com/watch?v=9JVNQzovcGQ). This is not surprising since these insects are associated with shade, humid conditions and litter.

*Caladenia flava* (cowslip orchid) flowers in late winter and early spring (Fig. 1E). Plants had shorter spikes with similar numbers of flowers per spike after fire (Fig. 7AB), but substantially more spikes formed per group (Figs. 4H, 7C, 9D). Pollination frequency was similar after fire (Fig. 8), so increased seed production resulted from greater flowering (Fig. 9D). Co-flowering plants required to support pollinators of this visually deceptive orchid were less common, but they were more visible after fire. Flower production was much lower the second year after fire, but also in unburnt areas, so fire promotion of flowering may last for several years but was most strongly regulated by climatic factors (Fig. S1). *Caladenia arenicola* (carousel spider orchid, Fig. 1J) pollination was significantly reduced (Fig. 8), but flowering was similar (Fig. 7) after fire.

*Diuris magnifica* (pansy orchid) is one of many very similar donkey orchids in the *D. corymbosa* complex (Fig. 1H). Pollination of this species and its close relatives is by visual deception of native bees that primarily visit Fabaceae flowers (Scaccabarozzi et al. 2020). Pansy orchid plants were substantially smaller after fire (Fig. 7A), but pollination marginally increased (Fig. 8), leading to a neutral response overall. Figure 9E suggests there is a stronger density effect on pollination in burnt areas, implying that pollinators may be less abundant after fire.

For *T. macrophylla* pollination fluctuated annually and was generally better in years with lower rainfall, presumably due warmer temperatures benefitting pollinators (Brundrett 2019). These have large blue flowers (Fig. 1L) primarily pollinated by native bees (Edens-Meier et al. 2013). As explained above, comparisons were limited by very low fire survival rates, but a few plants (8 spikes, 76 flowers) survived a small fire in 2023. These plants were shorter and had fewer flowers and seed than those in unburnt areas (average 9.5 vs. 15.5). Pollination was strongly regulated by plant density in unburnt areas (Fig. 9F).

*Pyrorchis nigricans* occurs across southern Australia and flowers in late winter. This very long-lived clonal species flowers almost exclusively after fires (Table 1). This orchid forms groups up to 30 m wide (average 3.4 m) primarily by clonal division (no seedlings were found). Reproduction is by multiple tubers on long droppers and large colonies would be hundreds or thousands of years old. *Pyrorchis nigricans* is relatively effective at attracting pollinators compared to other visually deceptive orchids in the same habitat and has comparatively large capsules (570 mm^3^), with up to 109 capsules in a group and an average of 14 (Brundrett 2019). Its flowers have no visible nectar or discernible scent so have an unknown type of attraction mechanism. A halictid bee has been observed visiting flowers in eastern Australia (Kuiter 2024).

Mass flowering of *P. nigricans* occurred after fires in 2019, when 14% of 128 spikes had capsules and in 2022, when thousands flowered and 8% of 2133 counted spikes formed capsules (Table 2, Fig. 1D). The higher rates of pollination of *P. nigricans* in 2019 may result from lower plant density in surveyed areas. There was a strong density effect on pollination in 2022, when 2/3 of capsules formed on the edge of groups (i.e. <20% of flowers formed 66% of seed). Single unburnt flowering plants were observed in 2020 and 2023 only (total of 8 flowers and 5 pods). Thus, flowering of this species is thousands of times greater post-fire. Some unburnt plants 5-10 m from the fire edge flowered in 2022 (34 plants). Thus, the flowering stimulus is air-borne and not heat and is likely to be ethylene gas (Dixon & Barrett 2003).

*Leptoceras menziesii* (rabbit orchid) also flowers primarily after fire (Fig. 1G) but sporadic flowering occurs at other times. In 2022 a group of 145 plants produced 54 spikes with 68 flowers that formed 44 capsules (65%), while in 2008 there were only 10 flowers on 6 spikes, with 30% seed set at a different location. This orchid also lacks nectar and has relatively high pollination rates.

Overall, orchid seed production was similar in burnt or long unburnt areas but was dominated by different species (Table 2). For 10 species that survived fire, seed production was largely unaffected (2 sp.), reduced (3 sp.), substantially increased (3 sp.), or induced by fire (2 sp.). Fire effects on pollination could not be assessed for seven species absent or rare in burnt areas. There were no major benefits of fire on reproduction for most orchids, which have adequate seed set to support local spread and population increases without fire (Brundrett 2019). However, *P. deformis,* produced much more seed after fire, and the pyrogenic species *P. nigricans* and *L. menziesii* rarely formed seed otherwise (Table 2).

### 4. Indirect fire effects

Longer-term effects of fire were determined by measuring the diversity and abundance of orchids relative to time since fire, so these results are independent of the data used to measure direct effects above. The fire in 2022 provided 18 ha for comparisons with long unburnt areas. Analysis of over 1000 orchid locations and 4.5 km of transects relative to fire history produced robust information for most species (Table 3, Fig. 11). These transects included a wide range of vegetation conditions and had limited overlap with location records (11%). Fire effects on orchids were analysed relative to time since fire (Fig. 12), since productivity was less well correlated with fire frequency (Figs. S2, S3). This would presumably be due to recovery of ecosystem functions and vegetation structure (Enright et al. 2011, Miller et al. 2025), as well as orchid recolonisation. Figure 11A shows that recent fire had a highly significant effect on the abundance of nine orchid species (P <0.05 from confidence intervals), and another seven species were absent or very rare after fire. Only *C. arenicola* was more abundant in recently burnt areas.

**Figure 11.**
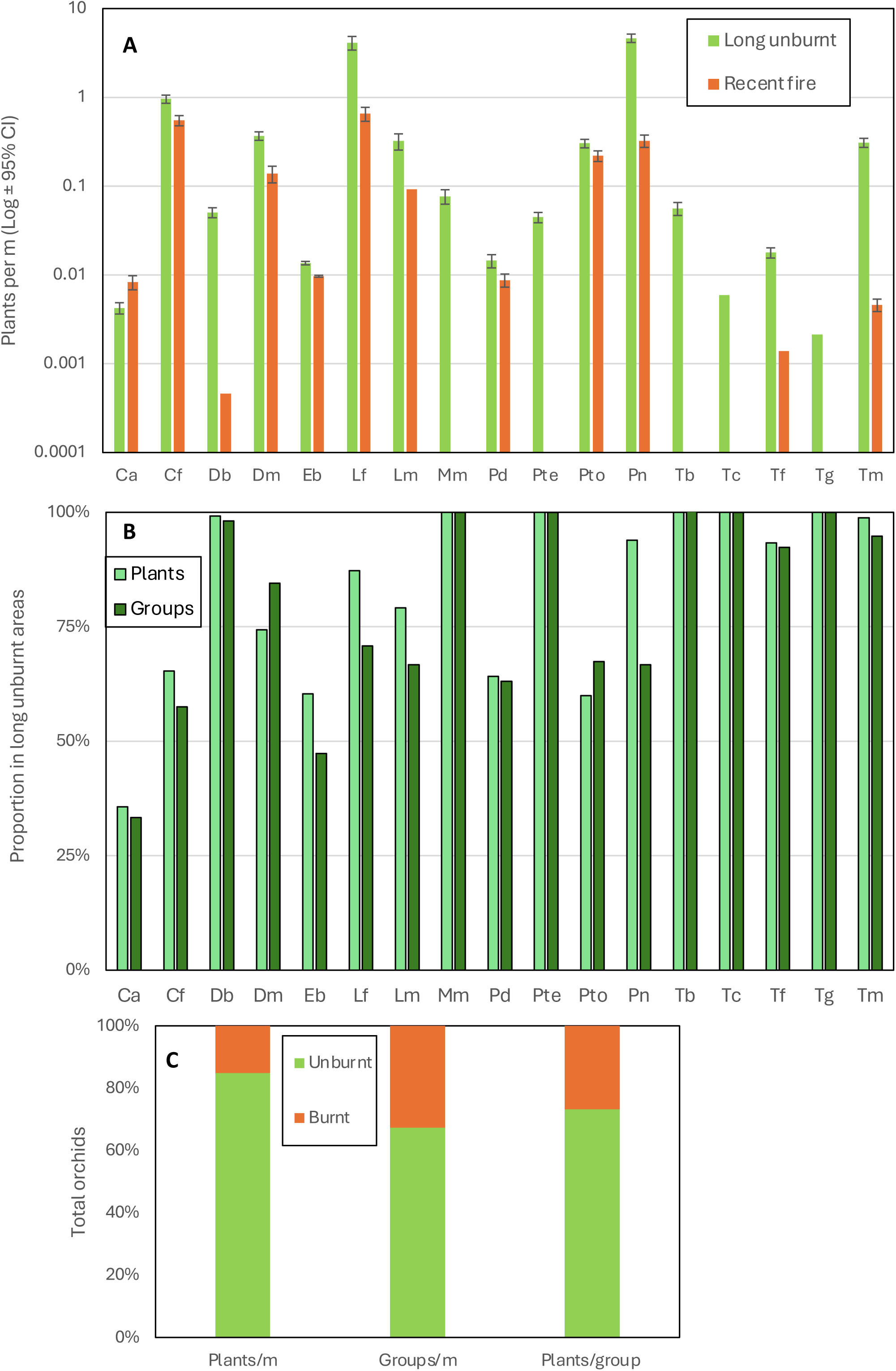
**A.** The relative abundance of 17 orchids in recently burnt (<3 years) or long unburnt (>20 years) areas on transects. **B**. The proportion of individuals or groups of orchids found in long unburnt areas on these transects. **C**. The relative proportions of all orchid plants or groups and average group sizes in recently burnt or unburnt areas.

**Figure 12.**
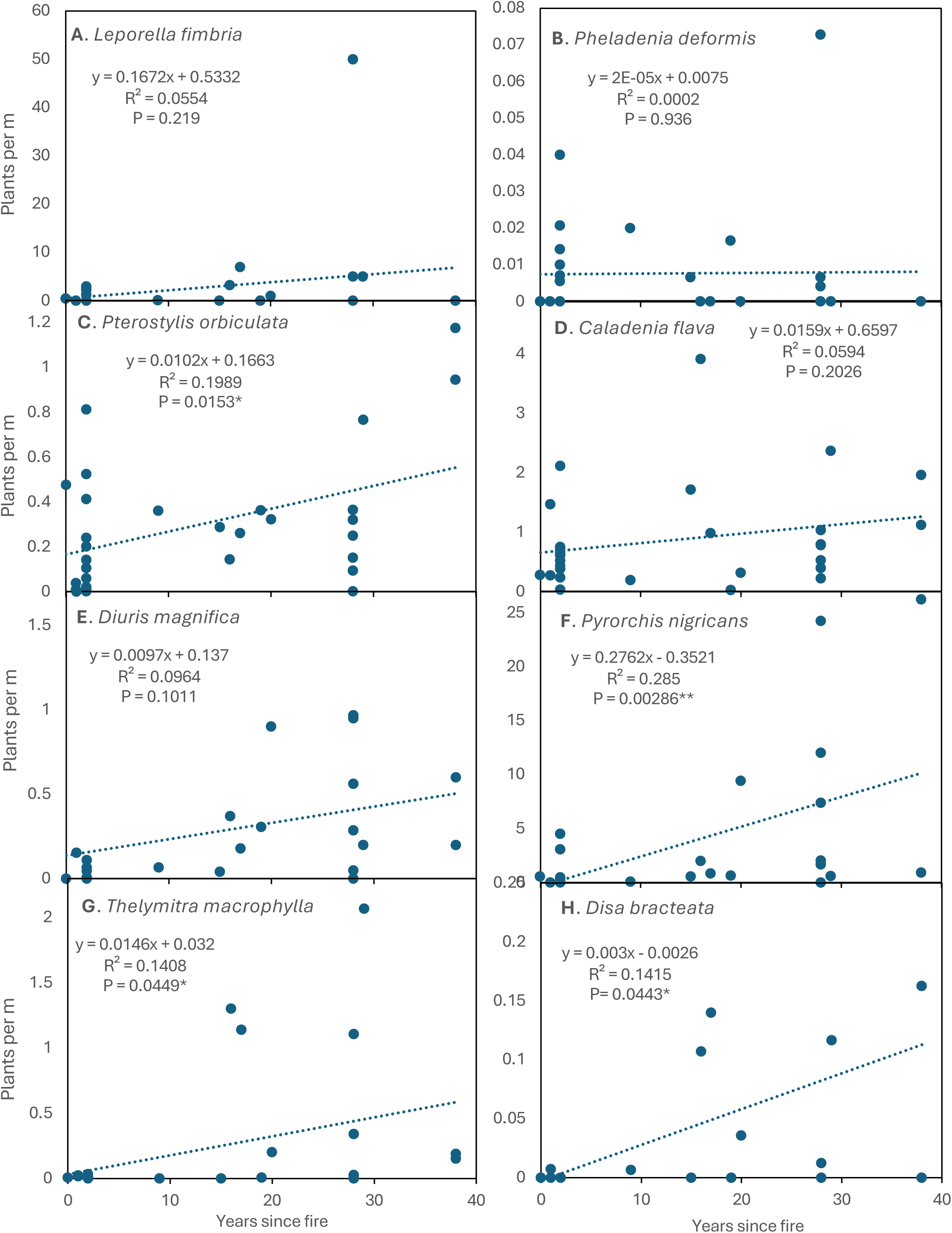
**A-H.** The abundance of eight common orchids (see labels) relative to time since fire using data from transects with regression equations and correlation probabilities.

As explained above, four small, relatively short-lived orchids that spread rapidly were extremely fire intolerant (*D. bracteata, M. media, P. ectypha* and *T. graminea*). They can rapidly colonise new areas by seed dispersal, provided substantial refugia exist nearby. However, this required at least 15 years peak abundance occurring several decades later (Fig. 12H, Table 3). In jarrah forest close relatives of *P. ectypha* were also most abundant several decades after fire (Cargill 2005) and began to colonise restoration areas 5 years post mining (Collins & Brundrett 2015).

Two very fire intolerant sun orchids (*T. macrophylla* and *T. graminea* - Fig. 1M) significantly increased in abundance after several decades (Figs. 11, 12G). Three other sun orchids had some degree of fire tolerance (*Thelymitra fuscolutea*) or lacked direct evidence of fire impacts (*Thelymitra benthamiana*, *Thelymitra crinita*). However, *T. fuscolutea* (Fig. 1Q) had a strong preference for growth in long unburnt areas (Fig. S2F), as also seems to be the case for *T. benthamiana* (Fig. 1N). *Thelymitra crinita* (Fig. 1K) was most abundant several decades after fire in the jarrah forest (Cargill 2005) and was absent from restored minesites but common in unmined forests nearby (Collins & Brundrett 2015). Seedlings of *T. benthamiana, T. crinita* and *T. macrophylla* were only found in older fire age areas (Fig. 4K, Table 3).

Two orchids survived and flowered well after fire but were more productive in long unburnt areas. *Pterostylis orbiculata* had a significant preference for shady areas under trees with leaf litter, where seedlings first appeared 3 years after fire (Figs. 4M, 11B, 12C). There also was evidence of increasing productivity with time since fire for *D. magnifica* (Fig. 12E). Orchids lacking a clear relationship with time since fire include *Caladenia flava* (Figs. 11, 12D) and *Elythranthera brunonis* (enamel orchid), which had similar abundance in burnt and unburnt areas (Figs. 1I, 11). *Caladenia arenicola* was found more often after fire (Fig. 11B), but had a flat response to time since fire (Figs. 12D, S2B). *Phelladenia deformis* also had a relatively flat fire age response (Fig 12A). It was the only orchid in this study likely to have substantially benefited from fires, due to strongly enhanced seed production, reproduction primarily from seed, relatively short lifespans and shade intolerance. Its bimodal fire age relationship mirrored the entire site (Figs. S2A, S2J).

In theory, the pyrogenic orchids *P. nigricans* and *L. menziesii* should have benefitted the most from fire, due to mass flowering. However, both were highly clonal and rarely recruited from seed, so these benefits could not be measured. Furthermore, *P. nigricans* was most abundant in long unburnt areas where it increased over time until some groups had thousands of leaves (Figs. 11, 12F). Gargill (2015) also categorised *P. nigricans* as an intermediate fire age species in jarrah forest. *Leptoceras menziesii* was also more abundant in long unburnt areas, as was *L. fimbria* (Figs. 11, 12A). Resolving fire age trends for tolerant species is complicated by inherently large variations in group sizes, but most benefited from long fire intervals. Overall, there were more orchids harmed by fire than benefited from it (Figs. 13, 14).

**Figure 13.**
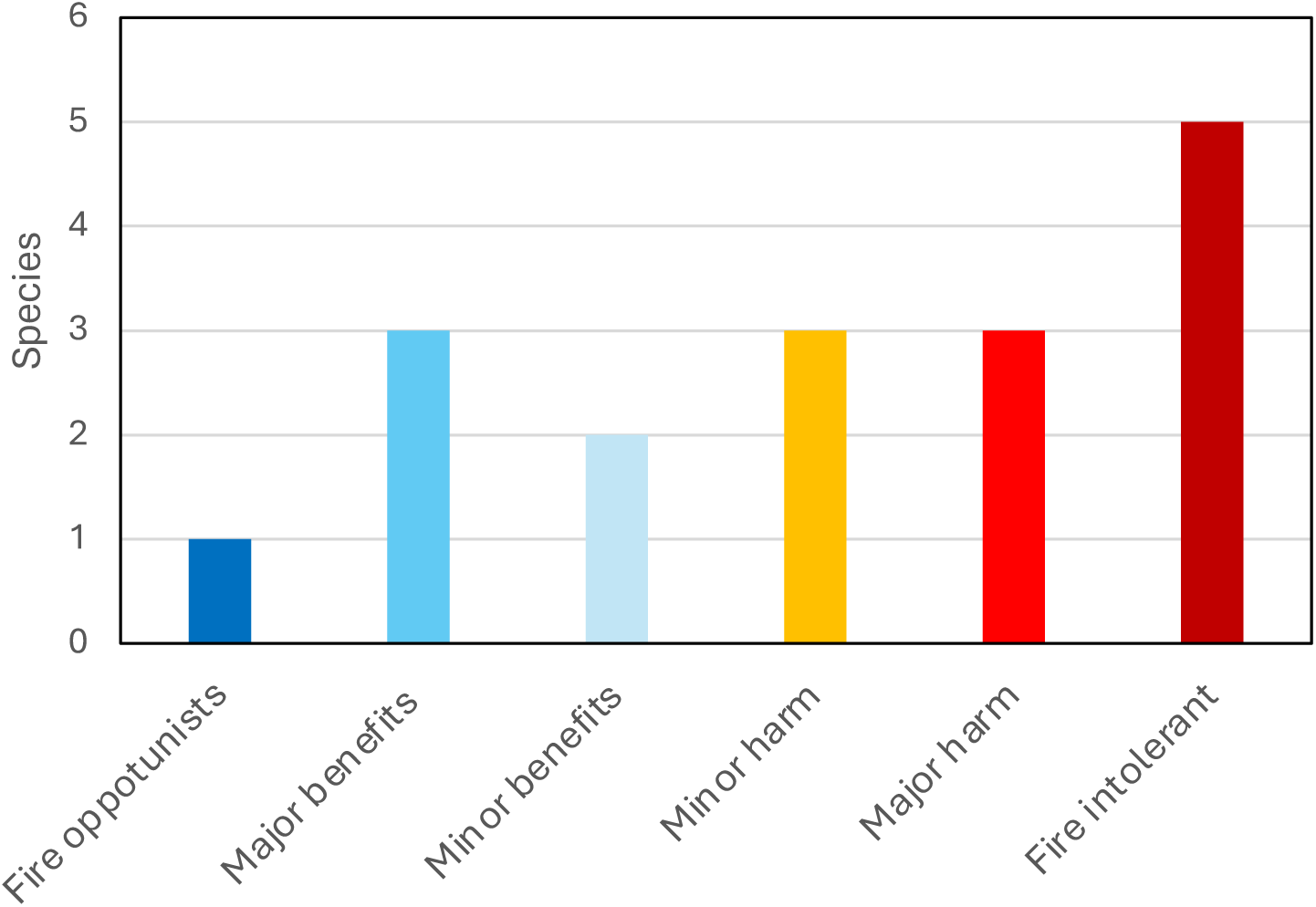
Total orchid density (**A**) and diversity (**B**) relative to time since fire on transects.

### 5. Comparing fire responses to other traits

Overall fire effects on orchid diversity and abundance were dramatic and statistically significant but differed substantially between orchid species so require classification in a consistent manner. The approach taken here was to develop fire tolerance scales based on criteria defined in Table 4. These indices are provided in Table 5 using data presented here (Tables 2, 6, Figs. 8-13). They include 10 categorical fire response indexes (Table 5A) with a common numeric scale (3 to -3), which were then combined into an overall Fire Response Index (FRI) for each orchid (-18 to 18). These FRI scores were used to assign orchids to tolerance categories (Fig. 15), as fire opportunist (13 to 18), major benefit (7 to 12), minor benefit (1 to 6), minor harm (0 to -6), major harm (-7 to -12) and fire intolerant (-13 to -18). Orchids identified as fire intolerant in Table 6 were normally lost to fire and increase in abundance several decades later, as illustrated by *T. macrophylla* and *D. bracteata* (Fig. 12G, H).

**Table 6.** Summary of reproductive and ecological traits and fire tolerance categories for orchids in the study (RAB - rare or absent after fire, FPF - fire promoted flowering).

| Orchid | Lifespan / aggregation | Primary means of reproduction | Seedlings or new plants | Disturbance response | Fire response category | FRI | FAST | Fire traits / responses | Ecological traits and fire responses | Colonisation potential in new habitats |
| --- | --- | --- | --- | --- | --- | --- | --- | --- | --- | --- |
| *Caladenia arenicola* | medium / low | seed | uncommon | weak positive | Minor benefits | 3 | 15 | fire neutral or positive | highly resilient, very deep tubers | colonised slowly (similar species) |
| *Caladenia flava* | long / medium | dropper tubers, seed | not seen | negative | Major benefits | 7 | 8 | FPF (weak) | more seed after fire | colonised slowly |
| *Disa bracteata* | short / low | abundant seed readily germinates | common | strong positive | Fire intolerant | -15 | 23 | RAB | exotic weed, very rare in burnt areas, RSS | colonised very rapidly then declines |
| *Diuris magnifica* | medium / medium | root tubers, seed | uncommon | weak positive | Minor harm | -5 | 15 | weak negative | highly resilient, very deep tubers, grows from tuber pieces | colonised rapidly (similar species) |
| *Elythranthera brunonis* | long / low | seed | not seen | negative | Minor harm | -2 | 9 | fire neutral | Short term decline after fire? | failed to return |
| *Leporella fimbria* | long / very high | dropper tubers, seed | not seen | negative | Minor benefits | 5 | 15 | FPF - strong | more seed after fire | failed to return |
| *Leptoceras menziesii* | long / very high | dropper tubers, seed | not seen | negative | Fire required | 13 | 10 | FPF - very strong | strongly promoted fire flowering | no data |
| *Microtis media* | short / low | seed | common | strong positive | Fire intolerant | -15 | 20 | RAB | weedy indigenous orchid, RSS | colonised rapidly, sporadic |
| *Pheladenia deformis* | medium / low | seed, tubers | uncommon | weak positive | Major benefits | 11 | 10 | FPF - strong | most seed produced after fire | spreading locally |
| *Pterostylis ectypha* | short / high | tubers, seed | uncommon | positive | Fire intolerant | -18 | 20 | RAB | absent in burnt areas, RSS | colonised rapidly (similar species) |
| *Pterostylis orbiculata* | medium / high | dropper tubers, seed | very common | weak positive | Major harm | -9 | 10 | stunted post-fire | first seedlings found 3 years after fire | colonised slowly (similar species) |
| *Pyrorchis nigricans* | very long / very high | dropper tubers, seed | not seen | negative | Major benefits | 10 | 25 | FPF - obligate & intolerant | near obligate fire flowering | very rare or never |
| *Thelymitra benthamiana* | short / low | seed | uncommon | negative | Major harm | -7 | 20 | no data | avoids burnt areas | colonised rapidly |
| *Thelymitra crinita* | medium / low | seed | uncommon | negative | Major harm | -8 | 20 | no data | many seedlings in 2022 | colonised very slowly, rare in restored areas |
| *Thelymitra fuscolutea* | medium / low | seed | uncommon | negative | Minor harm | -4 | 20 | limited data | avoids burnt areas | no data |
| *Thelymitra graminea* | short / low | seed | uncommon | negative | Fire intolerant | -18 | 25 | RAB | all groups lost to fire | no data |
| *Thelymitra macrophylla* | medium / medium | seed, tubers | common | negative | Fire intolerant | -12 | 20 | RAB | groups usually lost to fire | colonised rapidly, widespread |
| **Notes and data sources** | Table 5 | Table S3, Brundrett 2019, Pate & Dixon 1982 | Table 3 | local observations | from Fire Response Index, Table 5A | From Table 5 | FAST = Fire Age Safe Threshold | RAB - rare or absent post fire, FPF – fire-promoted flowering, RSS rapid spread by seed, |  | post-mining data from Grant & Koch 2003, Collins & Brundrett 2015 |

Data representing nine ecological traits expected to be related to fire responses (in Tables S1 to S5) were used to calculate indices with a unified scale (0-10), as provided in Table 5B. Comparisons between orchids revealed that fire responses were highly correlated with these traits, especially flowering, seed production, clonality, tuber depth, aggregation and lifespan (Table 7A, Fig. 16). One exception was tuber diameter which primarily reflected the size of orchids (Fig. 16B). Thus, fire susceptibility is deeply integrated into orchid ecology, especially traits that regulate dispersal and local persistence (Fig. 16). Comparisons of non-fire traits revealed that flower and seed production were inversely correlated with clonality, aggregation and lifespan indexes (Table 7A). The strongest relationship was an inverse correlation between lifespan and colonisation indexes (Fig. S5A, correlation coefficient = 0.95). Thus, orchids which reproduce primarily by seed are most effective at local dispersal, as would be expected. For similar reasons, indexes for aggregation and dispersion are also inversely correlated (Fig. S5B, correlation coefficient = 0.65). The dispersion index compares results from monitoring locations chosen to maximise orchid diversity, and unbiased sampling of orchids on transects (Fig 5), and was also correlated with the overall fire response index (Table 7A).

**Table 7A.** Correlation matrix for fire response indexes or traits for all orchid species.

|  | FAST | FRI | Flower<br>production<br>index | Seed<br>production<br>index | Colonisation<br>index | Clonality<br>index | Aggregation<br>index | Lifespan<br>index | Dispersion<br>index | Tuber radius<br>(mm) |
| --- | --- | --- | --- | --- | --- | --- | --- | --- | --- | --- |
| Fire age safe threshold (FAST) | 1 |  |  |  |  |  |  |  |  |  |
| Overall fire response (FRI) | -0.549 | 1 |  |  |  |  |  |  |  |  |
| Flower production index | 0.628 | -0.594 | 1 |  |  |  |  |  |  |  |
| Seed production index | 0.451 | -0.531 | 0.679 | 1 |  |  |  |  |  |  |
| Colonisation index | 0.328 | -0.741 | 0.606 | 0.419 | 1 |  |  |  |  |  |
| Clonality index | -0.316 | 0.453 | -0.619 | -0.416 | -0.706 | 1 |  |  |  |  |
| Aggregation index | -0.033 | 0.291 | -0.452 | -0.324 | -0.636 | 0.851 | 1 |  |  |  |
| Lifespan index | -0.401 | 0.764 | -0.596 | -0.415 | -0.950 | 0.631 | 0.576 | 1 |  |  |
| Dispersion index | -0.310 | 0.501 | -0.164 | -0.221 | -0.650 | 0.663 | 0.646 | 0.641 | 1 |  |
| Tuber radius (mm) | 0.234 | -0.158 | 0.260 | 0.270 | 0.105 | -0.081 | -0.126 | 0.062 | -0.010 | 1 |
| Tuber depth (mm) | -0.580 | 0.574 | -0.489 | -0.166 | -0.306 | 0.093 | 0.000 | 0.355 | 0.031 | -0.103 |

Correlation analysis in Table 7B revealed positive correlations between orchids that often co-occur due to shared habitat preferences (e.g. *C. flava, T. macrophylla* and *L. fimbria*). Very common orchids such as *T. macrophylla* and *P. nigricans* had strong positive correlations with total orchid density and diversity (Table 7B). In contrast, the abundance of *P. deformis* was inversely correlated with most other orchid species (and total orchid density and diversity), revealing disparate habitat preferences (Table 7B).

**Table 7B.** Species correlation matrix comparing relative abundance of common species.

|  | *Caladenia*<br>*flava* | *Disa*<br>*bracteata* | *Diuris*<br>*magnifica* | *Leporella*<br>*fimbria* | *Pterostylis*<br>*orbiculata* | *Pyrorchis*<br>*nigricans* | *Thelymitra*<br>*macrophylla* | *Pheladenia*<br>*deformis* | Total<br>plants | Fire age | All<br>orchids | Plants/<br>m |
| --- | --- | --- | --- | --- | --- | --- | --- | --- | --- | --- | --- | --- |
| *Caladenia flava* | 1 |  |  |  |  |  |  |  |  |  |  |  |
| *Disa bracteata* | 0.300 | 1 |  |  |  |  |  |  |  |  |  |  |
| *Diuris magnifica* | 0.135 | 0.312 | 1 |  |  |  |  |  |  |  |  |  |
| *Leporella fimbria* | -0.090 | -0.043 | -0.138 | 1 |  |  |  |  |  |  |  |  |
| *Pterostylis orbiculata* | 0.296 | 0.203 | -0.046 | -0.198 | 1 |  |  |  |  |  |  |  |
| *Pyrorchis nigricans* | 0.136 | 0.065 | -0.065 | 0.273 | 0.267 | 1 |  |  |  |  |  |  |
| *Thelymitra macrophylla* | 0.526 | 0.211 | 0.015 | 0.454 | 0.123 | 0.031 | 1 |  |  |  |  |  |
| *Pheladenia deformis* | -0.295 | -0.185 | -0.022 | -0.148 | -0.115 | -0.185 | -0.217 | 1 |  |  |  |  |
| Total plants | -0.009 | -0.031 | -0.027 | 0.931 | -0.102 | 0.466 | 0.376 | -0.121 | 1 |  |  |  |
| Fire age | 0.244 | 0.376 | 0.311 | 0.235 | 0.446 | 0.534 | 0.375 | 0.016 | 0.397 | 1 |  |  |
| All orchids | 0.494 | 0.016 | 0.256 | -0.113 | 0.190 | 0.176 | 0.428 | 0.091 | 0.037 | 0.469 | 1 |  |
| Plants/ m | 0.110 | 0.049 | -0.072 | 0.839 | 0.038 | 0.736 | 0.417 | -0.236 | 0.905 | 0.514 | 0.084 | 1 |
| Species / m | 0.463 | 0.249 | 0.019 | 0.029 | 0.345 | 0.465 | 0.564 | -0.306 | -0.008 | 0.503 | 0.446 | 0.335 |

Comparing orchid abundance and diversity to fire history revealed most species preferred long unburnt areas, including species that benefit from fire due to increased seed production (Figs. 11, 12, 13, 14). It also confirmed the rarity or absence of fire intolerant orchids in recently burnt areas (Figs. 5A, 14). Intersecting orchid locations with fire maps provided a Fire Age Safe Threshold (FAST) estimate for each orchid (Table 6). FAST values were difficult to precisely determine due to limited areas over 30 years post-fire (Fig. 12). Fire age is also highly correlated with overall orchid diversity and abundance (Fig. 13). The FAST values for orchids, which are based on long-term data, were highly correlated with their overall Fire Response Index (FRI), which primarily measures short term fire responses (Fig. 16H). Thus, two indices calculated from independent data sources provided similar fire tolerance classifications for orchids.

**Figure 14.**
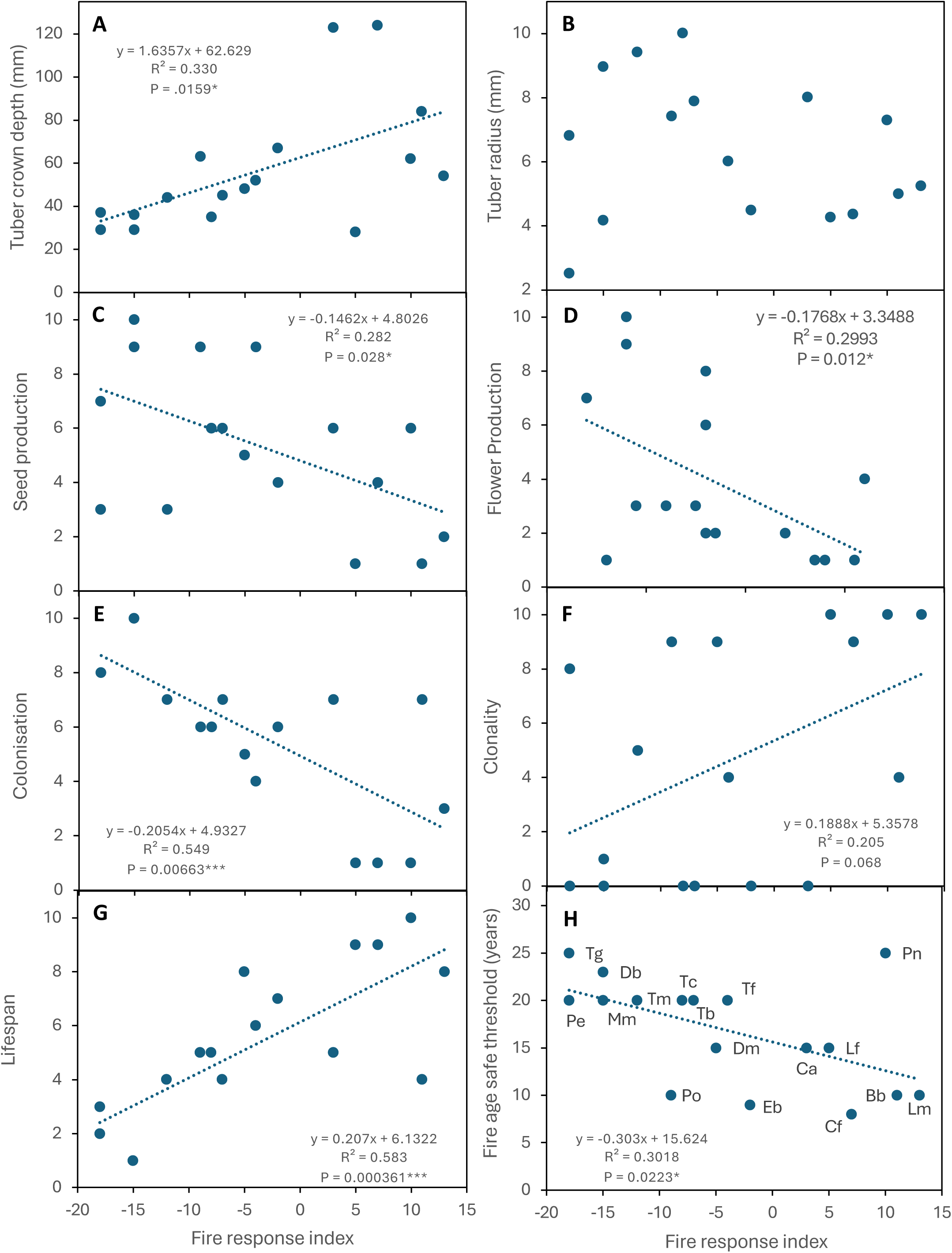
The average fire frequency and fire age where orchids occur (data from Table S4, name codes in Table 1). The average fire history for the entire site is also shown (red dot).

**Figure 15.**
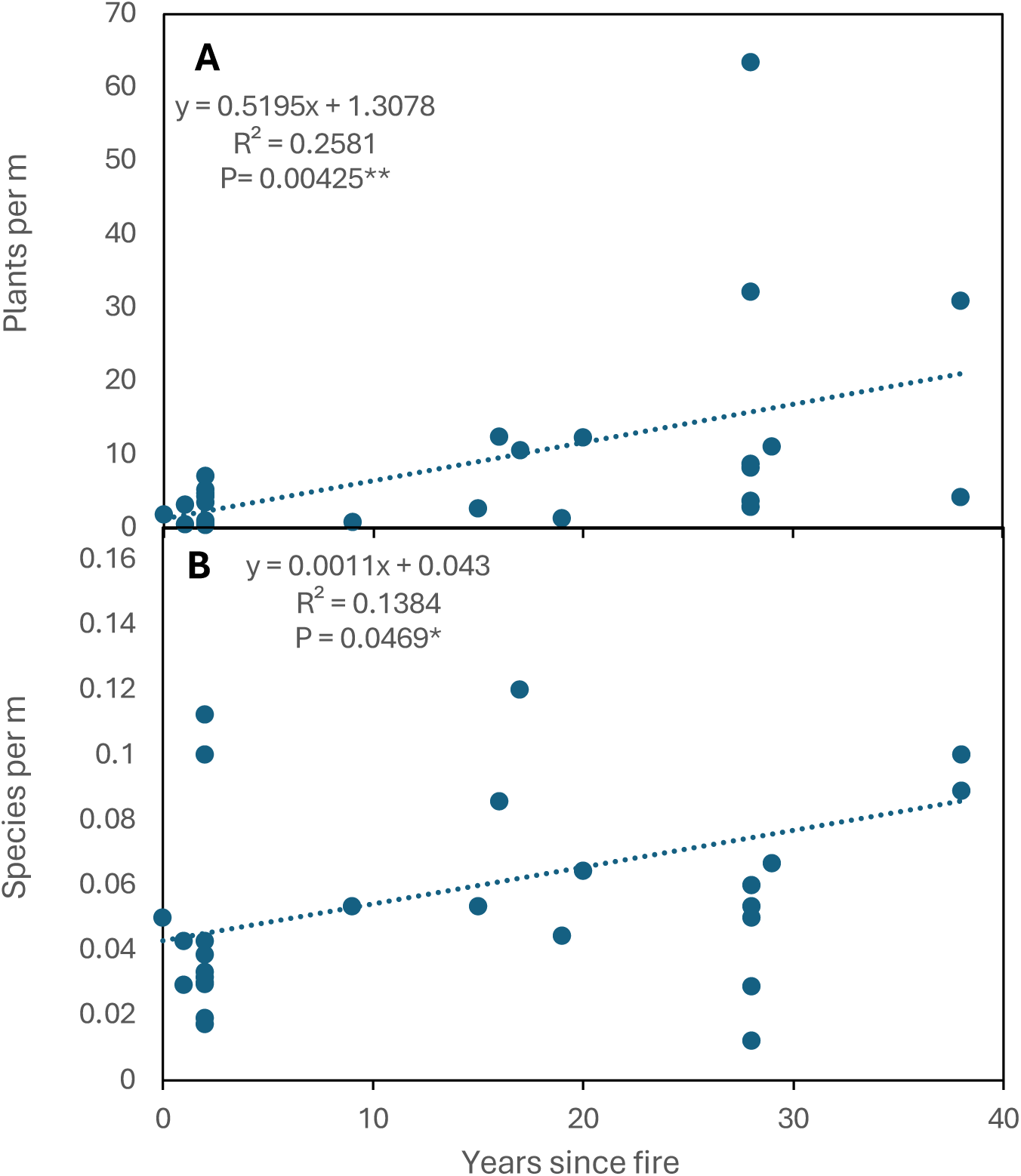
Overall fire tolerance rankings for all 17 orchid species.

**Figure 16.**
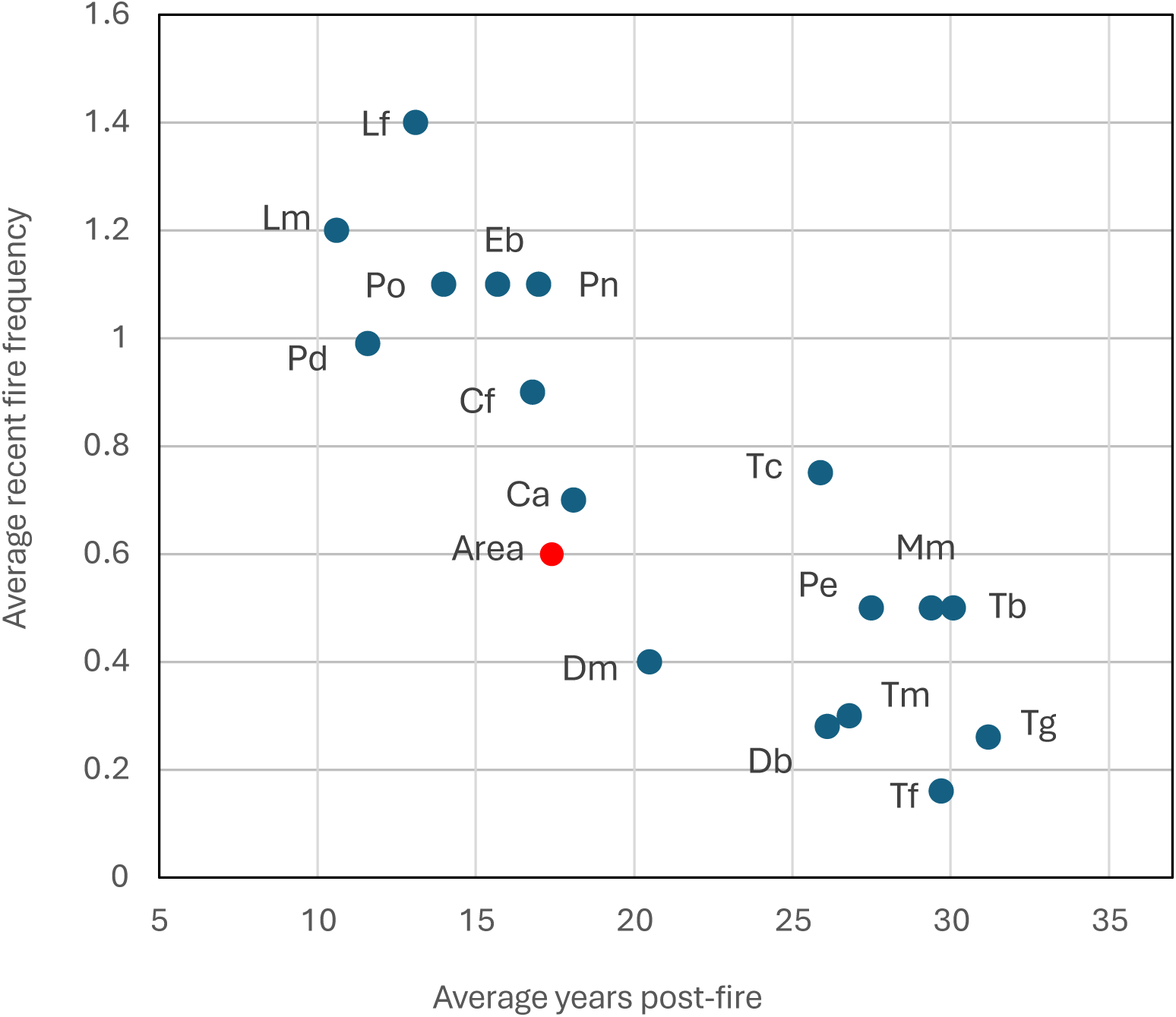
Correlations between tuber depth (**A**) or size (**B**) or key ecological trait indices, as labelled (**C-F**) with the overall fire response index for 17 orchid species.

Some orchids had similar abundance in recently burnt and long unburnt areas (Figs. 11, 12, 14). These included *E. brunonis, C. flava, P. deformis,* and *P. orbiculata*. *Caladenia arenicola* was the only orchid more common after fire, but this may be due to its preference for relatively open areas. However, even low FAST age orchids may require long unburnt areas for seedling establishment, where 86% of seedling nurseries were found (Table 3, Fig. 4). Thus, there are very few if any orchids that are fully fire promoted or neutral when relationships between habitat fire age and productivity or reproduction are considered.

### 6. Orchid fire ecology

Comparisons of orchid productivity, reproduction, diversity and abundance after fire to long unburnt areas revealed substantial variations in their fire tolerance (Box 1, Fig. 15), Fire response categories used here are based on those used by others (e.g. Jones 1988, Duncan 2012, Adams 2018), but were more precisely determined by using standardised measurements (Table 4). This revealed that orchids belonged to a continuum from obligate fire flowering to fully fire intolerant species, but it was difficult to assign some intermediate species to specific categories (see Figs. 14, 16H). There also were contradictions, such as *P. nigricans* which requires fire for flowering, but paradoxically is far more abundant in long unburnt areas where some colonies had thousands of leaves. This study highlights the importance of determining safe fire age thresholds (FAST) for orchids and their susceptibility to unseasonal fires (Box 1). Combined FAST data for orchid floras will also determine the time post-fire required to reach maximum orchid diversity and abundance in ecosystems.

##### Text Box 1. Fire response summary

1. Comparisons of recently burnt and long unburnt areas revealed clear short-term winners and losers after fire (Table 6, Fig. 15):

a. Four orchids had major fire benefits and two had minor benefits.
b. Six species experienced minor or major harm from fire due to reduced productivity.
c. Five orchids rarely survived fire or were entirely absent from recently burnt areas.
2. Three orchids had pyrogenic (fire-induced) flowering (Lamont & Downes 2011, Bower 2019, Brundrett 2021). Two of these were very long-lived clonal species that rarely reproduced by seed, so benefits from seed production may be minimal, and *P. nigricans* was far more abundant in long unburnt areas. The third species (*P. deformis)* formed most of its seed after fire, is relatively short lived and propagates primarily by seed. This orchid may function as a fire opportunist but was also common in long unburnt areas (Fig. 11).
3. Orchids intolerant of fire include two sun orchids *T. macrophylla* (89% fire mortality) and *T. graminea* (100% mortality). Three other fire intolerant species were small, relatively short-lived and spread rapidly due to abundant seed production (*M. media, D. bracteata* and *P. ectypha*). These orchids required several decades to recolonise burnt areas (Table 3).
4. At least five orchids that were normally resilient to fire could be lost to unseasonal fires, especially in autumn (Table 5A), as was also found in eastern Australia (Jasinge et al. 2018, Thomsen et al. 2024).
5. Overall, eight species were relatively common in recently burnt areas, while nine species were found much more often (4 sp.) or only (5 sp.) in long unburnt areas (Figs. 11, 14).
6. The total FRI score for all orchids studied here is biased to the negative (-3.8) since more species had negative overall fire effects even if they survive well and flower more after fire (Table 5A).

In this study, orchid diversity and abundance increased consistently with time since fire, which was more important than the total number of fires in an area since their impacts diminished over time (Fig. 13). This study demonstrates the importance of long-term orchid monitoring and data from the same areas before and after fire. There are very few other studies of orchid productivity relative to time since fire (Duncan 2012).

The ecology and biology of terrestrial orchids in an ancient biodiversity hotspot is highly correlated with their fire responses. As explained above, these relationships were revealed using trait indexes, many of which are correlated with each other (Table 7A). In particular, fire tolerant orchids were more likely to be aggregated, have long lifespans, reproduce clonally and have relatively deep tubers (Figs. 16, S5). In contrast, fire intolerant orchids tended to have greater flower and seed production, reproduce primarily by seed, spread more rapidly, have shallower tubers and shorter lifespans. However, these orchids would have less need for fire resilience because they are relatively mobile, provided dispersal from long unburnt habitats nearby can occur.

Fire intolerant orchids sometime survived in fire areas, presumably due to fire patchiness and intensity differences that were not revealed by fire mapping. For example, some *T. macrophylla* plants survived a winter fuel reduction fire after substantial leaf scorching, but this species usually failed to survive intensive summer fires when it was dormant. This discrepancy may be associated with relatively shallow tubers or a preference for leaf litter and soils high in organic matter (see below). Fire impacts on habitat quality may explain why *T. macrophylla* population viability had not recovered five years after a managed winter fire. *Leporella fimbria, P. nigricans* and some *Thelymitra* or *Pterostylis* species initiate shoot growth in mid-autumn resulting in tuber depletion and vulnerable new shoots. These shoots are held just below the soil surface until there is sufficient rain, so can be killed by fires at these times (see Fig. 4G). These orchids will not become resilient to fire until tubers are replaced over winter. The importance of fire intensity and seasonality in determining orchid fire responses require further investigation.

Orchid fire traits may vary across the ranges of widespread or variable species. For example, *P. nigricans*, which has an Australia wide distribution, is more likely to flower without fire in habitats which rarely burn (Brundrett 2014). Other widespread orchids that may have variably fire traits include *P. orbiculata, D. magnifica* and *T. macrophylla*, which are indistinct taxa in species complexes (Brundrett 2014). Intraspecific variability in fire traits is well documented in other SWAFR plants (Brundrett 2021).

Fire response data presented here are from one banksia-jarrah woodland site in an urban setting with an exceptionally high fire frequency. However, these data agree with results of studies of the same or closely rated species in other SWAFR habitats (Table 6). These include 9 of the same species in a jarrah forest fire study (Cargill 2005) and 14 of them from post-mining recolonisation of jarrah forest (Collins & Brundrett 2015). Adams (2018) found similar relationships between fire age and orchid diversity in eastern Australia. Orchids with traits likely to make them fire intolerant include 37% of SWAFR species (Brundrett 2021). Taken together there is strong evidence to support fire tolerance variability (measured by FAST and FRI) as a keystone of Australian orchid ecology.

In this study, fire substantially modified habitat suitability for orchids by increased light from canopy loss, reduced organic matter and increased dominance of major environmental weeds, especially veldt grass (*Ehrharta calycina*). Some orchids were less common in vegetation with many weeds, but others persisted well in these areas. These factors were addressed here by including similar areas of weedy and non-weedy vegetation on transects. However, interactions between fire history, vegetation condition and the relative cover of native plants or weeds require further study.

The ability to recolonise or spread within areas by wind dispersed seed is very important for orchids after fire. Many SWAFR orchids have highly specific relationships with insect pollinators and mycorrhizal fungi, so their habitats must also be suitable for their fungal and insect symbionts (see introduction). These relationships are more one-sided (exploitative) than for other SWAFR plants (Brundrett 2021). For example, 84% of SWAFR orchids have deceptive pollination syndromes that exploit insects (Brundrett et al. 2024). These insects include native bees for visually deceptive orchids such as *C. flava* and *D. magnifica*, or wasps, ants or fungus gnats for sexually deceptive orchids such as *C. arenicola, L. fimbria* and *Pterostylis* species (Scaccabarozzi et al. 2018, 2020, Kuiter 2024). Pollination is also affected by pollinator decline in isolated urban habitats where these insects must compete with abundant feral honeybees that rarely transfer pollinia (Scaccabarozzi et al. 2024). Despite all this, most pollination systems studied here were highly resilient to fire or promoted by it, provided plants survived. This has also been observed in other fire-prone biomes (Duncan 2012, Brown & York 2016; Tsiftsis et al. 2025).

Orchids are parasites of fungi, at least for part of their life cycles, while the majority of other SWAFR plants have mutualistic mycorrhizal associations (Brundrett 2021). Seed baiting trials have shown that orchid fungi were associated with soil organic matter (Brundrett et al. 2003), and their germination was limited to small patches of soil in banksia woodland (Batty et al. 2001). Thus, orchid habitats are partly defined by soil organic matter, leaf litter and shade required by their mycorrhizal fungi, which are all severely impacted by fire. This would explain why orchid seedlings were primarily found in long unburnt areas. Fungi germinating orchid seeds were also detected more often in forest than in restored minesites in the jarrah forest, where they were associated with litter accumulation in rip lines (Collins et al. 2007). Unfortunately, little is known about the ecology of orchid pollinators or fungi, including their capacity to survive intense fires or recolonise habitats.

As summarised above, the fire intolerance of orchid species is determined by interacting processes including (i) fire-induced mortality, (ii) reduced seed production, (iii) dispersal or recruitment failure, (iv) reduced soil quality and (v) habitat conditions, such as loss of shade and increased weed cover. These factors work in concert, but their relative importance and the time required for recovery of each are unknown. In contrast, fire tolerant orchids avoid mortality and are less impacted by these factors, but their abundance often decreases after fire. The FAST status of orchids is likely to be regulated by both orchid and site-specific factors, but only the former was measured here. For example, the time required for orchids to recover could be longer in isolated patches of suitable habitat due to limited seed dispersal. New orchid species occasionally arrive in isolated urban areas such as Warwick Bushland, where five new species were recently discovered, but often did not persist. These “vagrant” orchids include *Caladenia falcata*, a common species further east, which temporarily appeared in five separate locations.

### 7. Management implications

As explained above, sustainable management of orchid populations and habitats requires a thorough understanding of life history traits regulating their persistence and dispersal in ecosystems, and how they interact with fire history and climate extremes. Key trait data required to manage orchids includes their lifespans, turnover, seed production, dispersal efficiency and clonal reproduction, but are lacking for most Australian species. Maintaining local populations of orchids also requires an understanding of essential resources including symbiotic organisms such as mycorrhizal fungi and pollinating insects. Very long-lived clonal species can be resilient in the short term, but this makes it challenging to determine impacts of fire on their sustainability. Short lived orchids with weed-like life history traits require conditions that favour seed production and seedling establishment and good connectivity between natural areas to persist. They may also benefit from intermediate levels of disturbance such as grazing, or fire (Coates et al. 2006, Hutchings 2010). Thus, both short and long-lived orchids require FAST and FRI data for effective management. Strong relationships between fire history and orchid diversity suggests that intolerant orchid diversity could be used to estimate fire age in ecosystems, but this requires validation.

Here, the optimal fire age for orchid diversity was found to be at least 30 years in banksia woodland. Similar correlations between fire age and orchid diversity also occur in SWAFR forests (Cargill 2005). Fuel reduction fires in autumn, winter, or spring are especially damaging for orchids, since they initiate shoot growth in autumn or winter and form tubers several months later (Brundrett 2014, Thomsen et al. 2024). Spring fires are also very damaging because most orchids are flowering or forming seed at that time (Brundrett 2014). Unfortunately, both autumn and spring fuel reduction fires are commonly applied to SWAFR orchid habitats (Bradshaw et al. 2018). Other major fire-induced changes to orchid habitats included substantially increased weed cover, canopy reduction and more understory competition due to pyrogenic seed germination and resprouting of native plants and weeds. It may be possible to promote recovery of burnt orchid habitats by augmenting soils with organic materials, weed management, artificial shade, etc., but this is likely to be a slow process.

Fire tolerance indicators such as FAST and FRI concepts should be relevant to other plants and organisms – especially fungi and small animals involved in recycling organic matter which is consumed by fire. Research to establish such data should be a very high priority in ecosystems where fire impacts are increasing. The relative impacts of fuel reduction fires relative to very hot summer fires on FAST times is another critical knowledge gap, as is the impacts of unseasonal fires which impact orchids when they are most vulnerable and could lead to local extinctions.

As explained in Box 2, Fire Age Safe Thresholds are an essential tool for sustainable management of orchids in Australian ecosystems. This approach is critically important for managing rare species, where core habitat areas must be excluded from managed fires. The Wheatbelt Orchid Rescue project mapped core habitat areas, for five extremely rare orchids (Brundrett 2016), but 39 other threatened SWAFR orchids and 72 “priority flora” species require further investigation (florabase.dbca.wa.gov.au, 28-10-24). Setting maximum burn areas and frequencies for managed fires and monitoring their biodiversity impacts is required to maintain orchid diversity (Box 2). Habitat management should aim to maintain a mosaic of fire ages, with fuel reduction fires used to protect long unburnt vegetation in core habitat areas for fire intolerant organisms. These refugia would provide nurseries for recruitment and seed production for dispersal (Box 2). These approaches would also protect focal points for tourism and protect local biodiversity hotspots such as exposed granite, sheoak or wandoo woodlands, damplands, peat soils, organic soils, shady areas, habitat trees and rotten logs.

##### Text Box 2. Orchid fire management

1. Orchids have diverse fire responses which require separate analysis of direct and indirect effects on their survival and productivity, as well as knowledge of local fire history to be understood.
2. Ecological and biological trait indexes (FRI) and fire age safe thresholds (FAST) provide valuable tools for habitat management for orchids and other fire intolerant organisms. They also provide a framework for understanding the local distribution, turnover and abundance of species.
3. Some of the most beautiful and iconic SWAFR orchid species are fire intolerant (e.g. Fig. 17).
4. Fire ages longer than three decades are optimal for most orchids and longer times may be required by some species. Few orchids were more productive with shorter intervals, even if they tolerate them.
5. When fire response data is lacking a precautionary approach is required, using fire ages longer than FAST values for similar species or resulting in maximum diversity and abundance in similar habitats.
6. Arson is the main cause of fire in urban Perth and results in major damage to local ecosystems, but lacks effective management. Severe droughts and groundwater decline magnify fire impacts in these areas but lack sufficient recognition or effective management.
7. Effective weed management in banksia woodland substantially reduces impacts on vegetation due to fire. Control of grazing by rabbits or other animals is also important.
8. Some orchids are relatively short-lived dynamic species efficiently spread by wind-dispersed seed, while others are much more static, forming large groups hundreds or thousands of years old. Impacts of fire are much more concerning for the latter.
9. Nature reserve management should aim to maximise the diversity of orchids, and other fire intolerant flora and fauna.
10. Orchid seedling recruitment primarily occurs in long unburnt areas with litter accumulation and relatively few weeds. Landscapes should be managed to protect these nurseries.
11. Understanding ecological impacts from fuel reduction activities requires evidence that benefits out way their costs, especially if fire frequency is already unsustainable due to arson and climate change.
12. Orchids have a major role in attracting visitors to natural areas, but changes to fire management may be required to ensure this continues. These iconic species can promote the importance of healthy ecosystems, especially in urban areas (e.g. friendsofwarwickbushland.com/walk-trails/jarrah-trail).
13. Measuring orchid diversity is a relatively easy and effective way to monitor fire impacts on ecosystem health and can utilise existing or new data obtained by volunteers. Studies in urban areas provide essential knowledge for conserving less accessible rare orchids.
14. Effective management of rare species requires enforceable fire management goals and protocols and effective monitoring that triggers remedial actions. An alarming case study is provided by the critically endangered Queen of Sheba orchid (*Thelymitra variegata*) which has less than 25 remaining wild plants. A key population declined substantially after a fuel reduction fire (MCB unpublished).

### 8. Conclusions

This study presents a comprehensive analysis of orchid fire ecology in an urban nature reserve with a very complex six-decade fire history. Combining fire maps with orchid distribution and flowering data revealed a continuum from very fire tolerant to fully intolerant species. Fire traits allowed orchids to be classified into distinct risk categories defined by tolerance indices (FRI) and safe fire intervals (FAST), that were based on independent data. Even though pyrogenic flowering can lead to amazing orchid displays in the SWAFR, the harm caused by fire to orchids is substantially greater than its benefits overall. Furthermore, some species that require fire for seed production are most productive in long unburnt areas. This paradox likely resulted because evolution of orchid fire traits was driven by very infrequent intense fires in the SWAFR, following Gondwanan origins of plant fire traits in the Cretaceous (Lamont & He 2012). Consequently, orchid habitats require substantial areas with a fire age of several decades to maintain sustainable populations. This study includes representatives of 10 genera including the four most important in Australia, so its conclusions are likely to be relevant in other habitats. More research on orchid fire ecology is urgently required in Australia and globally, using tools such as those introduced here.

**Figure. 17.**
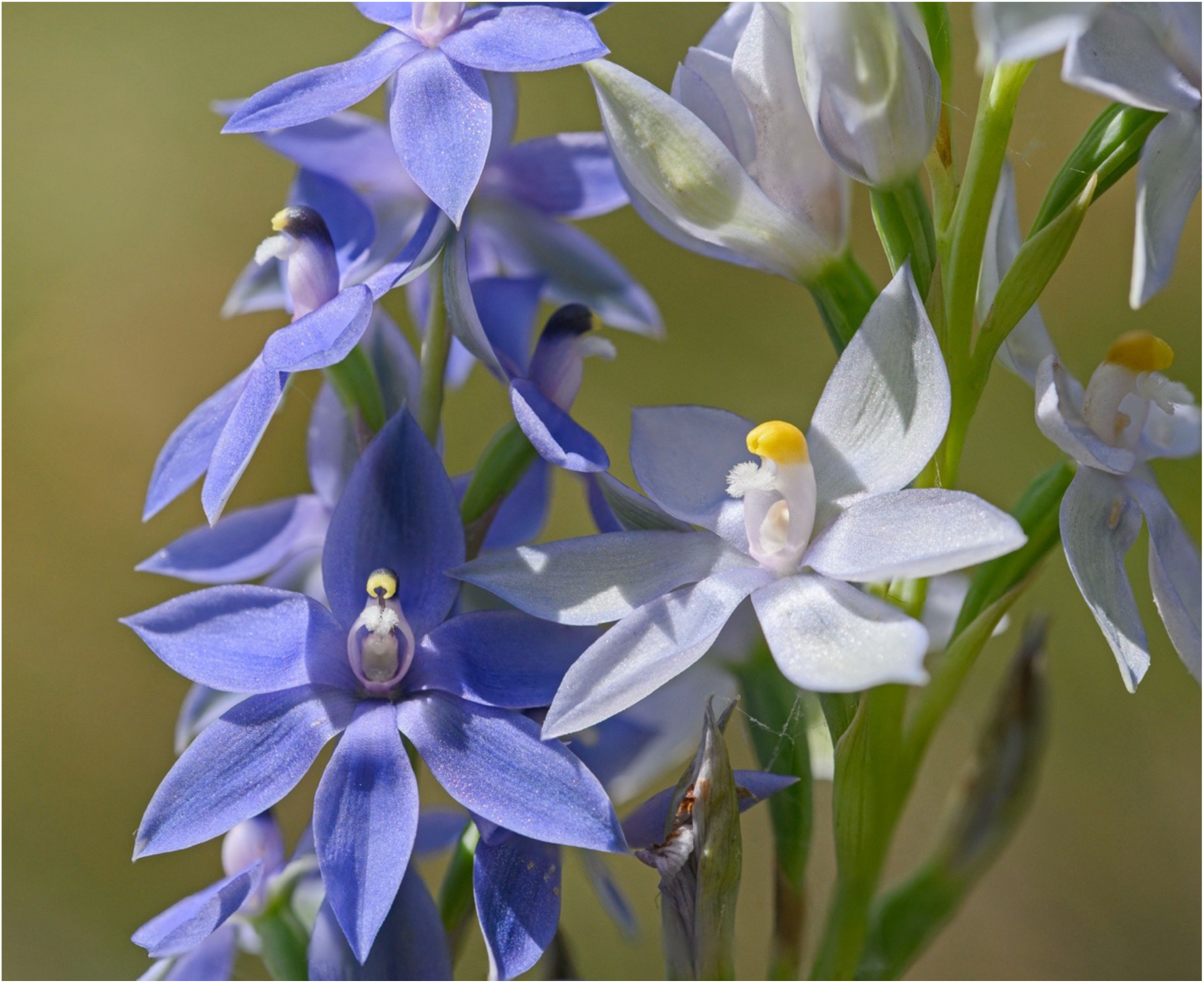
The blue sun orchid (*Thelymitra macrophylla*) requires areas without fire for 15 to 30 or more years to thrive. This orchid attracts many visitors to Warwick Bushland and is a focus of education activities (friendsofwarwickbushland.com/walk-trails/jt020).

While, some orchids are remarkably resilient to fire, others are highly vulnerable, so are threatened in Western Australia by increasing frequencies of managed fires, arson and climate driven extreme fire events (Bradshaw et al. 2018, Bergstrom et al. 2021, Udy et al. 2024). Managing orchids during this man-made crisis requires an understanding of relationships between time since fire and orchid population sizes, as well as their mechanisms for recolonisation of burnt areas. The intensity, frequency and timing of fires is also important, and unseasonal fires have major impacts on some species. Orchid fire ecology is closely integrated with other key aspects of their ecology and biology, such as tuber depth, clonality, dispersion and disturbance tolerance. These orchid traits can be measured using indexes that help to predict future risks from changing fire regimes. Quantification of orchid life history traits also revealed complex mechanisms at the heart of Australian orchid ecology requiring further analysis.

Results presented here suggest that mixed fire ages in landscapes are essential to maintain orchid diversity, with long unburnt areas the most important overall (Box 2). Thus, fire intolerant orchids require large nature reserves that include substantial areas of long unburnt vegetation. Fires managed to reduce fuel can also drastically impacts orchid habitats, because these “fuels” include organic matter required by their symbiotic fungi. Under current fire regimes orchid diversity is threatened in many habitats in Western Australia, and even common orchids, which are a key resource of sustainable ecotourism, are likely to experience widespread declines.

## Acknowledgements

Special thanks to Karen Clarke for patience, joining me on many walks through the study site to discover local biodiversity. I am also very grateful to many others for help locating orchids, especially Tim Hodgkins and other Friends of Warwick Bushland, as well as City of Joondalup staff. Daniella Scaccabarozzi commented on the manuscript.

## Data Availability Statement

Data needed to evaluate the conclusions in the paper are presented in the Supporting Information.

---

**Figure S1.**
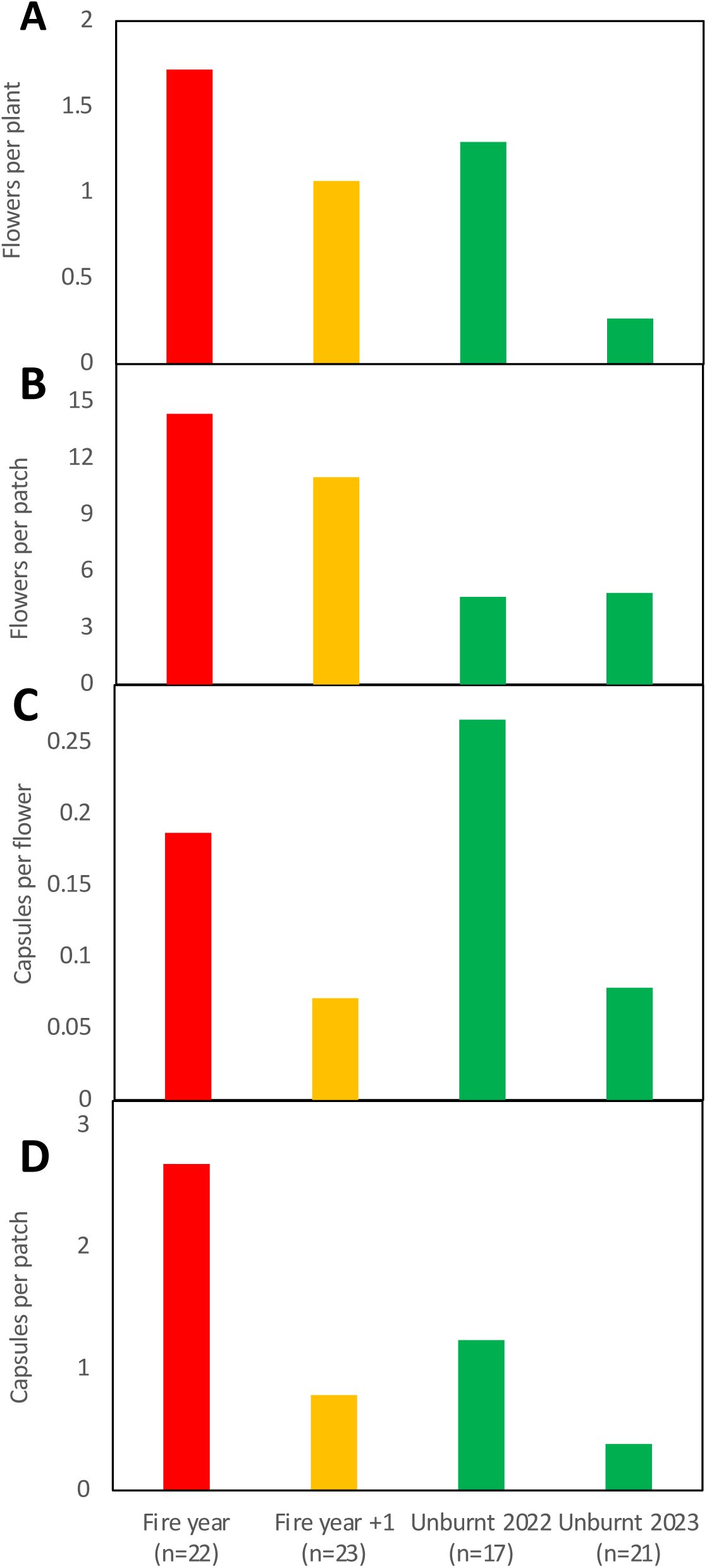
*Caladenia flava* flowering trends over two years. Graphs compare flowering (**A-B**) or pollination (**C-D**) in burnt or unburnt areas on the same two subsequent years.

**Figure S2.**
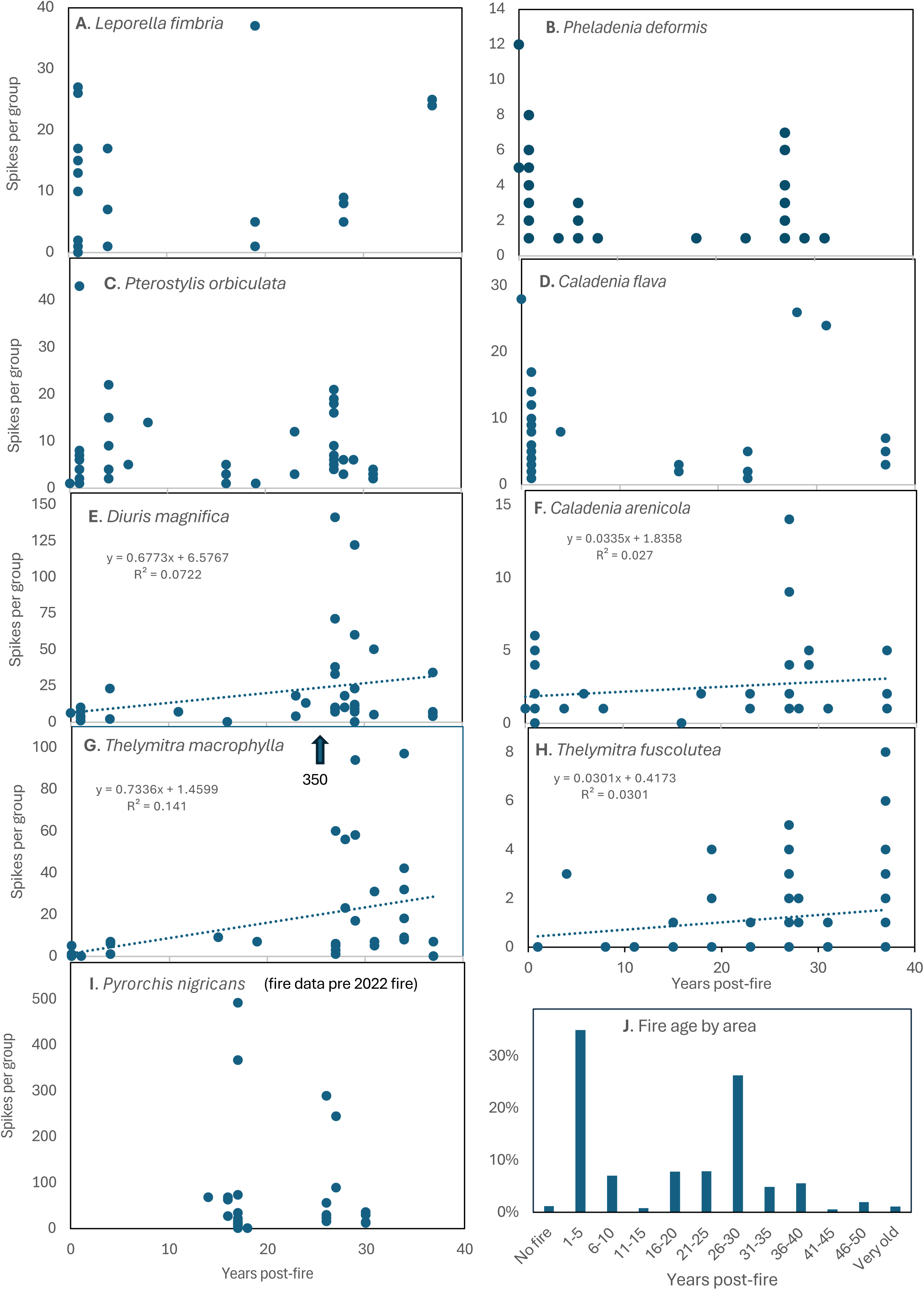
The productivity of groups of eight common orchids relative to time since fire measured by the number of flowering plants in monitored areas (**A-I**). The overall frequency distribution of fires by area is also shown (**J**).

**Figure S3.**
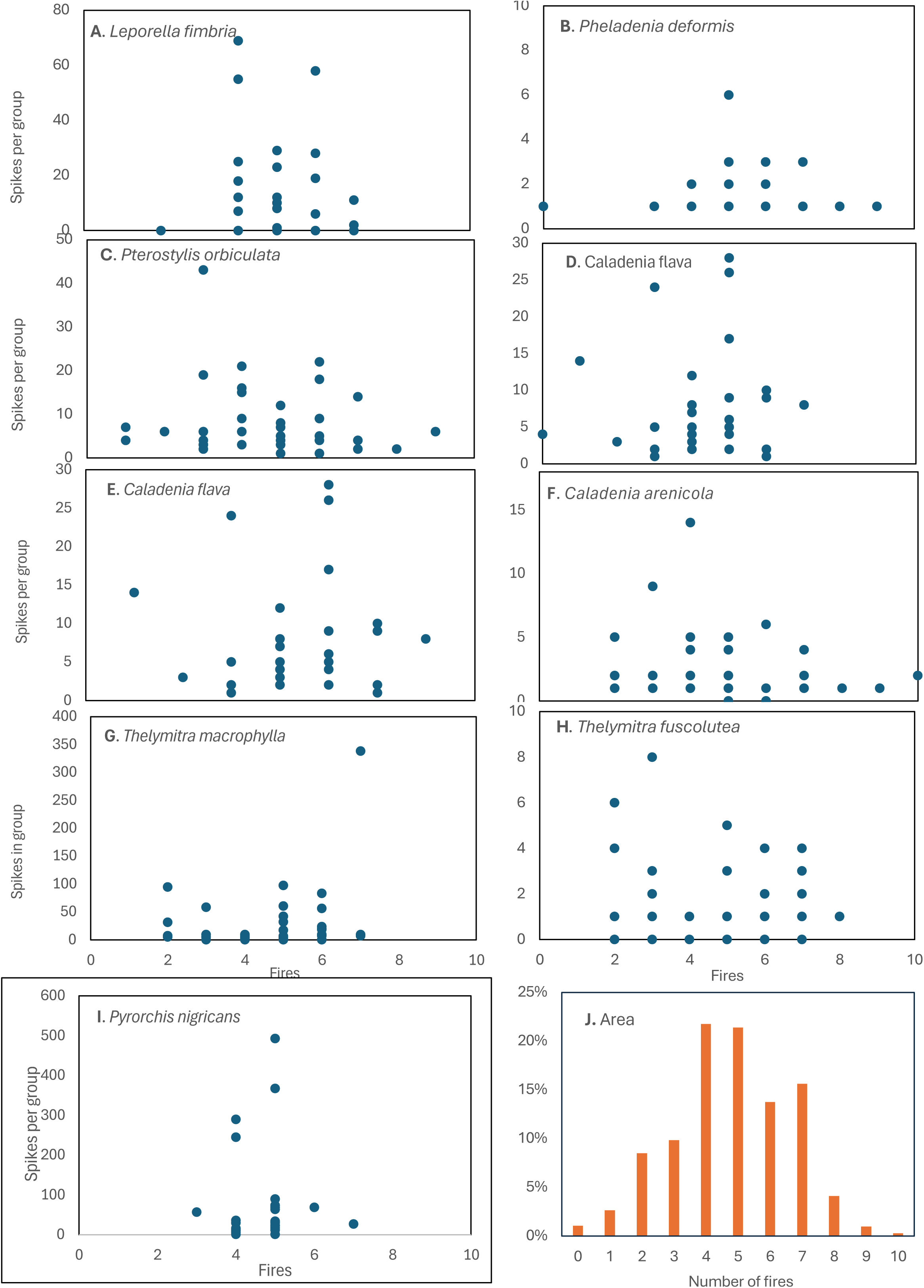
Relationships between frequency of all fires and flowering productivity of eight locally common orchids (**A-I**). The overall frequency distribution of fires by area is also shown (**J**).

**Figure S4.**
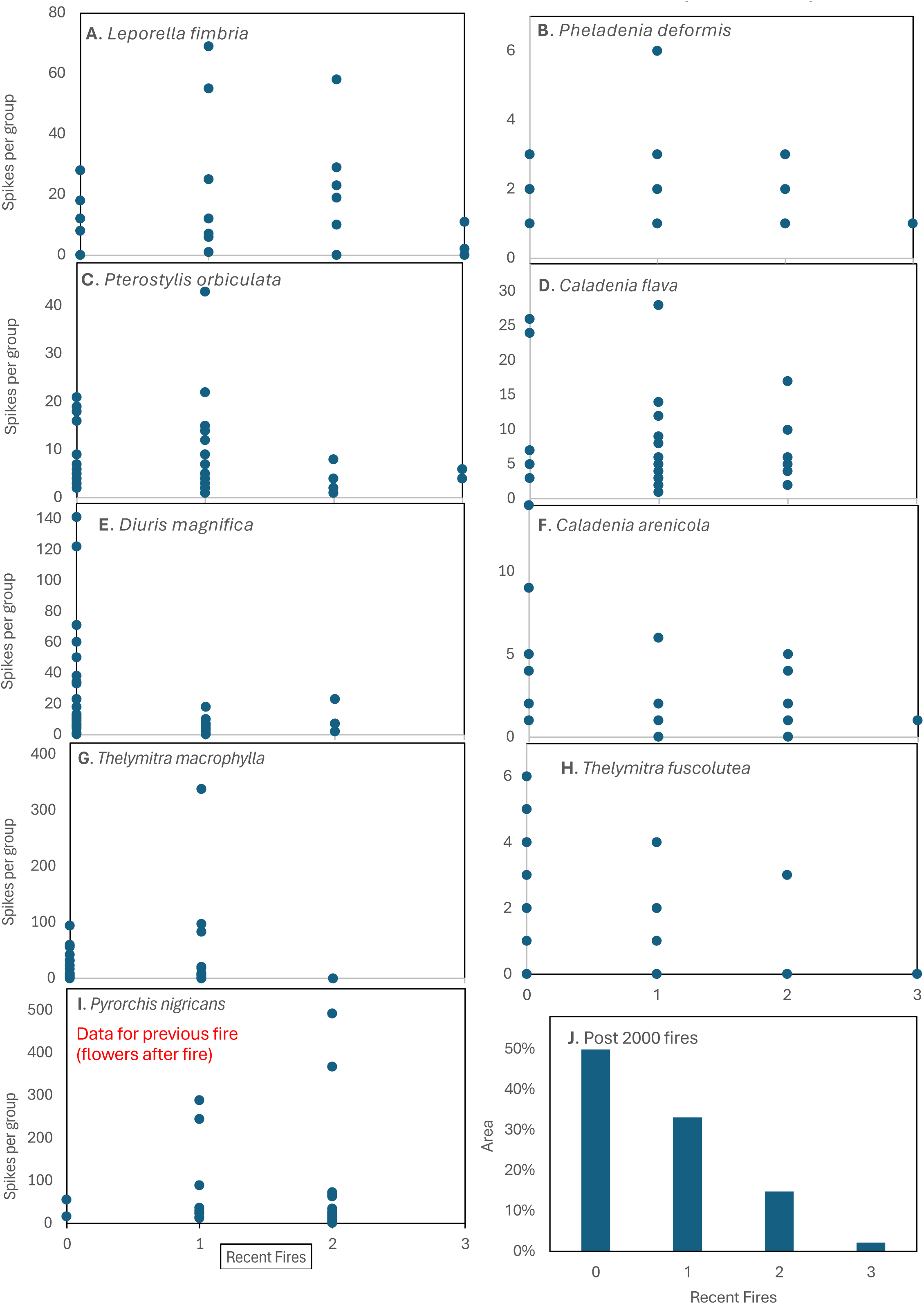
The productivity of groups of nine common orchids (**A-I**) relative to the frequency of recent fires (2000-2023). The overall frequency distribution of fires by area is also shown (**J**).

**Figure S5.**
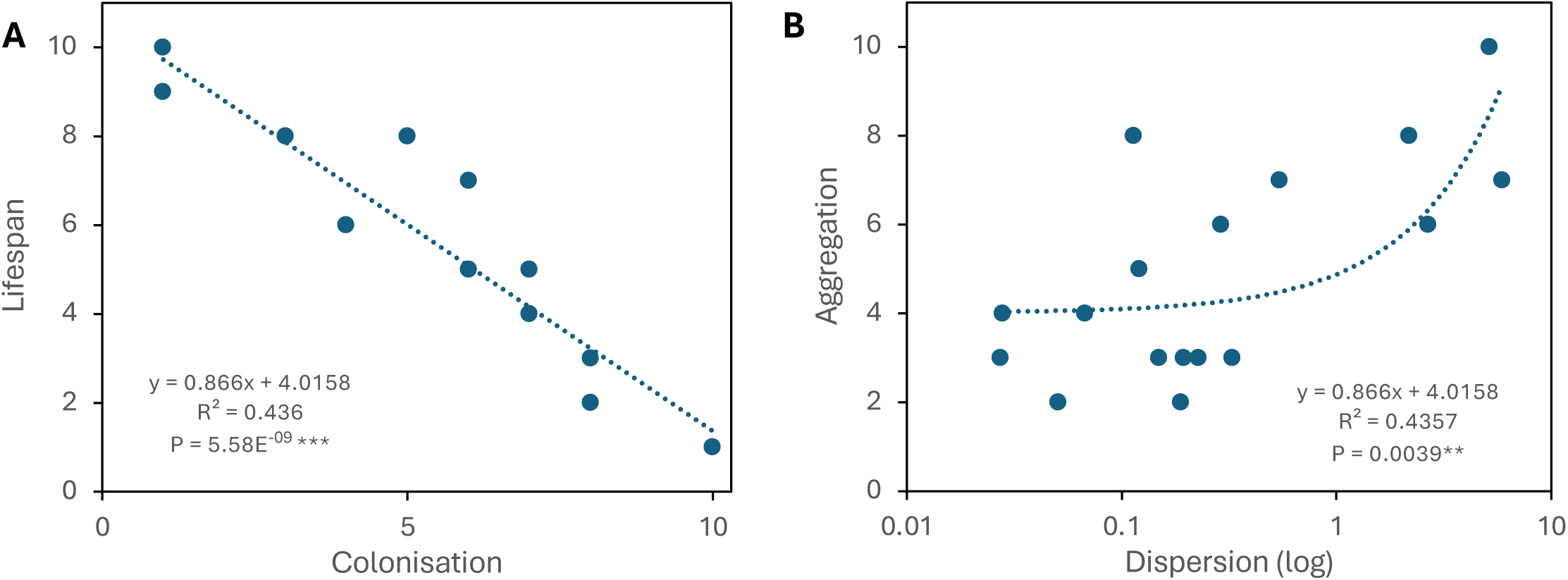
Additional trait correlation graphs. **A.** Estimated lifespan and colonisation capacity. **B**. Aggregation and dispersion.

